# Characterization of prohibitins as novel target to block asexual and sexual stage growth of *Plasmodium falciparum*

**DOI:** 10.1101/2022.06.02.494630

**Authors:** Monika Saini, Che Julius Ngwa, Manisha Marothia, Vandana, Kailash C. Pandey, Soumya Pati, Anand Ranganathan, Gabriele Pradel, Shailja Singh

**Author notes:** **Correspondence:** Shailja Singh.

## Abstract

Prohibitins (PHBs) are highly conserved pleiotropic proteins as they have been shown to mediate key cellular functions. Here, we characterize PHBs encoding putative genes of *Plasmodium falciparum* by exploiting different orthologous models. We demonstrated that *Pf*PHB1 (PF3D7_0829200) and *Pf*PHB2 (PF3D7_1014700) are expressed in asexual and sexual blood stages of the parasite. Immunostaining indicated these proteins as mitochondrial residents as they were found to be localized as punctate foci. We further validated *Pf*PHBs as organellar proteins residing in the *Plasmodium* mitochondria, where they interact with each other. Functional characterization was done in *Saccharomyces cerevisiae* orthologous model by expressing *Pf*PHB1 and *Pf*PHB2 in cells harboring respective mutants. The *Pf*PHBs functionally complemented the yeast PHB1 and PHB2 mutants, where the proteins were found to be involved in stabilizing the mitochondrial DNA, retaining mitochondrial integrity and rescuing yeast cell growth. Further, Rocaglamide (Roc-A), a known inhibitor of PHBs and anti-cancerous agent, was tested against *Pf*PHBs and as an antimalarial. Roc-A treatment retarded the growth of PHB1, PHB2, and ethidium bromide petite yeast mutants. Moreover, Roc-A inhibited growth of yeast PHBs mutants that were functionally complemented with *Pf*PHBs, validating *P. falciparum* PHBs as one of the molecular targets for Roc-A. Roc-A treatment led to growth inhibition of artemisinin-sensitive (3D7), artemisinin-resistant (R539T) and chloroquine-resistant (RKL-9) parasites in nanomolar ranges. The compound was able to retard gametocyte growth with significant morphological aberrations. Based on our findings, we propose the presence of functional mitochondrial *Pf*PHB1 and *Pf*PHB2 in *P. falciparum* and their druggability to block parasite growth.

## INTRODUCTION

Malaria has been a severe infection since decades killing up to 627, 000 people in 2020 alone. The number of malaria infected cases and deaths increased in the year 2020 due to the COVID-19 pandemic where it deeply impacted the malaria prevention, diagnosis and treatment services primarily in Sub-Saharan African regions (1). Accounted for causing most of the reported human malaria cases, *Plasmodium falciparum* holds global importance. However, a vast majority of its proteins are yet to be characterized experimentally to understand the crucial cellular pathways leading to pathogenesis of the organism (2). Mitochondria being one of the essential organelles of the parasite, has been serving as a molecular target for various antimalarial drugs. In this study, we have focused our attention in delineating the characterization and functional significance of *Pf*PHBs in *Plasmodium* mitochondria in addition to proposing these as drug targets. We have attempted to characterize these two putative members of superfamily of proteins called <u>S</u>tomatin/<u>P</u>rohibitin/<u>F</u>lotillin/<u>H</u>fiKC (SPFH) in the *P. falciparum*, focusing primarily on *Pf*PHB1 (PF3D7_0829200) and *Pf*PHB2 (PF3D7_1014700) and elaborating their functional significance in both asexual and sexual blood stages of the parasite. PHB1 and PHB2 are two highly homologous subunits of the mitochondrial PHB complex (3–6). PHB1 and PHB2 are interdependent on the protein level, and the loss of one simultaneously leads to the loss of the other (7, 8). Both proteins consist of an evolutionarily conserved PHB (Band-7/SPFH) domain that is similar to that of lipid raft associated proteins, and a C-terminal coiled-coil domain which is involved in protein–protein interactions, including the interaction between PHB1 and PHB2 (9). The location of PHBs has been a controversial topic as they form functional complexes in the inner mitochondrial membrane apart from being regarded as nuclear and membrane proteins (10–13). However, evidences suggest that PHBs have more critical roles within the mitochondria and mediate crucial mitochondrial processes, including protein processing, cristae morphogenesis, mitochondrial DNA organization, stress tolerance, and respiratory chain functions (13–16). At the inner mitochondrial membrane, PHB1 and PHB2 are known to form a 1-2 MDa heterodimeric ring-like supra-macromolecular structure (diameter 20-25 nm) and maintain the mitochondrial stability via this PHB1/PHB2 interaction under metabolic stress (10, 11, 13, 15). Recently, a study based on homology modelling and yeast two-hybrid analysis showed that putative *Plasmodium* PHBs, interact with each other suggesting that they could form super-complexes of heterodimers in *Plasmodium*, the functional form required for optimum mitochondrial function (17). As evident, PHBs play a crucial part in maintaining the mitochondrial homeostasis supported by various studies as mentioned above.

In line with emphasizing mitochondrial functions, stage-specific development of parasite requires enormous amount of energy, fueled by the power house organelle-mitochondria (18). This double membrane organelle of the parasite is an important engagement site for essential metabolic pathways such as electron transport chain, tricarboxylic acid (TCA) cycle and many more (2, 19, 20). The membrane of mitochondria itself is a commencing point and linking bridge for signalling these metabolic pathways. Therefore, the trafficking around mitochondrial membrane is highly regulated by specialized integral membrane protein, ensuring the parasite’s survival (20). Any compromise in regard to the membrane integrity is detrimental for the protein trafficking, eventually shutting down the parasite development (19). PHBs, are one such integral membrane protein responsible for maintaining mitochondrial protein homeostasis essential for parasite development (21). Despite these diverse biological roles, the establishment of PHBs as drug targets remains underestimated (22).

Based on the available literature, Rocaglamides (Roc-A) have been shown to bind to PHBs in cancer cells and inhibiting the downstream functional pathways (22, 23). Roc-A, cyclopenta[*b*]benzofurans, belonging to a class of molecules called flavaglines are secondary metabolites isolated from the plant genus *Aglaia* of the family *Meliaceae*. Till date, more than 100 naturally occurring Rocaglamide-like compounds have been isolated displaying anti-cancer properties at nanomolar concentrations (24–26). The primary mode of action by Rocaglamides on tumor growth inhibition was considered to be due to the protein synthesis inhibition without any effect on DNA or RNA synthesis (23). Interestingly, rocaglamide has been used as an anticancer drug to treat enterovirus 71 neuro-pathogenesis via studying the role of prohibitins in the virus biology (27). Another recent study highlighted the rocaglate CR-1-31B in disrupting the association of *P. falciparum* eIF4A (PfeIF4A) with RNA (28). In line with this, we explored the antimalarial activity of Roc-A against *P. falciparum* infection by targeting both *Pf*PHB1 and *Pf*PHB2, and proposing these proteins to be effective drug targets through drug repurposing.

With this background knowledge, we aimed to characterize the putative *Pf*PHB1 and *Pf*PHB2 proteins in the blood and transmission stages of *P. falciparum* for a better understanding of the functional aspect of these proteins. We demonstrated that these proteins are expressed throughout the asexual and sexual blood stages of *P. falciparum* development. Interestingly, both *Pf*PHB1 and *Pf*PHB2 were found to be expressed and focused as a punctate structure in gametocyte stages, corresponding to the presence of a single mitochondrion in the sexual stages of *P. falciparum*. Further findings confirmed *Pf*PHBs to be integral membrane proteins where they form the functional complex by interacting with each other. Molecular and biophysical approaches confirmed the interaction of *Pf*PHBs with Roc-A, where the drug binds to the target proteins and efficiently inhibits the parasite growth with IC_50_ in nanomolar ranges. Functional complementation in *S. cerevisiae* PHB mutants rescued the yeast growth significantly. Additionally, *S. cerevisiae* PHB mutants were rescued by complementation with *Pf*PHBs in petite-mutants (with disrupted mitochondrial DNA). Hence, our finding deepens the available knowledge of PHB proteins in *P. falciparum* and presents *Pf*PHB1 and *Pf*PHB2 as essential mitochondrial proteins required for maintaining mitochondrial integrity. Moreover, this study could be beneficial in employing PHBs as target-based drug designing approaches in *Plasmodium*.

## MATERIAL AND METHODS

### Gene identifiers

The following PlasmoDB gene identifiers are assigned to the *P. falciparum* genes and proteins analyzed in this study: PHB-1 [PlasmoDB: PF3D7_0829200]; PHB-2 [PlasmoDB: PF3D7_1014700]; STOML [PlasmoDB: PF3D7_ 0318100]; PHBL [PlasmoDB: PF3D7_0416600]; Pfs230 [PlasmoDB: PF3D7_0209000]; MSP-1 [PlasmoDB: PF3D7_0930300]; Pf39 [PlasmoDB: PF3D7_1108600]; PfAldolase [PlasmoDB: PF3D7_1444800]; Pfs25 [PlasmoDB: PF3D7_1031000]; H_4_Kac_4_ [(tetra)-acetyl histone H_4_ K_5, 8, 12, 16_ac]; AMA1 [PlasmoDB: PF3D7_1133400]; PfCCp2 [PlasmoDB: PF3D7_1455800]; GAP45 [PlasmoDB: PF3D7_1222700], NapL [PlasmoDB: PF3D7_1203700]

### Bioinformatics

Protein sequences of *Pf*PHB1 (PF3D7_0829200) and *Pf*PHB2 (PF3D7_1014700) were retrieved from PlasmoDB database. Ab-initio models of *Pf*PHB1 and *Pf*PHB2 were generated through I-TASSER server and validated (29). Further, the resulting models were subjected to structural refinement by using ModRefiner (30). The refined three-dimensional models generated were considered as the final structures.

### Cultivation of *P. falciparum* parasite culture

*P. falciparum* parasite strains (3D7, NF54, R539T, and RKL-9) were cultured in RPMI 1640 media (Gibco, USA) supplemented with 50 mg/L hypoxanthine (Sigma-Aldrich, USA), 2 gm/L sodium bicarbonate (Sigma-Aldrich, USA), and 10 mg/L gentamicin (Gibco, USA) supplemented either with 5 gm/L AlbuMax I (for 3D7, R539T and RKL-9; Gibco, USA) or 10% heat inactivated human A+ serum (for NF54) using human erythrocytes, under mixed gas environment (5% O_2_, 5% CO_2_ and 90% N_2_) following the procedures of Trager and Jensen (31). To induce gametocytogenesis in NF54, lysed RBCs were added to cultures maintained at high parasitaemia. The cultures were supplemented with 50 mM N-acetyl glucosamine (GlcNac) for approximately 5 days once stage I gametocytes appear in order to kill the asexual blood stages in the gametocytes. The gametocytes were then maintained in normal culture medium without GlcNac until immature (stage II -IV) or mature stage V gametocytes were enriched by Percoll gradient centrifugation. Activated gametocytes were obtained after incubating Percoll-enriched mature gametocytes with 100 µM xanthurenic acid (XA) dissolved in 1% 0.5 M NH_4_OH/ddH_2_O for 15 minutes at room temperature (32, 33)

### Transcript expression analysis of SPFH superfamily

To determine the transcript expression of *spfh* superfamily, asexual blood stages (rings, trophozoites and schizonts) of synchronized NF54 parasite cultures were harvested, whilst, immature and mature gametocytes (Gc) were enriched via Percoll gradient purification, and the activated gametocytes were collected at 30 minutes post activation. Total RNA was isolated from respective stages using the TRIzol reagent (Invitrogen, USA) according to the manufacturer’s protocol. Genomic DNA contamination was removed by treating the isolated RNA samples with RNase-free DNase I (Qiagen), followed by phenol/chloroform extraction and ethanol precipitation. Photometric analysis revealed A260/280 ratios > 2.1. The RNA concentration of the samples was determined using a ND-1000 (NanoDrop Technologies, Thermo Fisher Scientific, USA) and subjected to complementary DNA (cDNA) synthesis. 1 µg of total RNA from each sample was reverse transcribed into cDNA using the SuperScript IV First-Strand Synthesis System (Invitrogen, USA), following the manufacturer’s instructions. In order to determine if the gametocytes samples were not contaminated with asexual blood stages, the cDNA was tested by diagnostic RT-PCR using primers specific for the gene encoding the apical membrane antigen (*Pfama-1*, 189 bp, asexual blood stage contamination) and for gametocyte-specificity using primers specific for the gene encoding the LCCL-domain protein *Pfccp2* (198 bp). Transcripts for *Pfphb1* (196 bp), *Pfpbh2* (155 bp), *Pfstoml* (229 bp) and *Pfphbl* (213 bp) were amplified in 25-35 cycles using respective primers (**Table S1**). Amplification of *Pfaldolase* (378 bp) was used as a loading control and to investigate potential contamination with gDNA in the negative control lacking reverse transcriptase. PCR products were separated by 1.2% agarose gel electrophoresis. The gels were subjected to densitometry to calculate the relative intensity of each band.

### Molecular cloning, purification of *Pf*PHB1 and *Pf*PHB2 recombinant proteins

*P. falciparum* phb1 and phb2 nucleotide sequences were selected for recombinant protein generation as a fusion protein with pGEX-4T 1 vector (Ambersham Biosciencces, UK; with a cleavable Glutathione-S-Transferase (GST) tag at the N-terminus) and pMAL^TM^c5X (New England Biolabs, USA; with an N-terminal maltose-binding protein (MBP) tag). DNA fragments encoding specific regions of the genes (*Pf*PHB1: 1-819 bp for pGEX-4T 1 and 115-819 bp for pMAL^TM^c5X; *Pf*PHB2: 1-915 bp for pGEX-4T 1 and 61-915 bp for pMAL^TM^c5X) were PCR amplified from genomic DNA (*Pf*3D7) using gene-specific primer pairs (**Table S1**) with Phusion™ High-Fidelity DNA Polymerase (Thermo Scientific, US). The amplified DNA fragment was purified with QIAquick Gel Extraction Kit (Qiagen) using the manufacturer’s protocol. Purified inserts and expression vectors pGEX-4T 1 and pMAL^TM^c5X were digested with BamHI/SalI (for pGEX-4T 1 constructs) and NotI/PstI (for pMAL^TM^c5X constructs) restriction enzymes (New England Biolabs, UK) and were ligated over night at 16 °C using T4 DNA ligase (New England Biolabs, UK). The ligation mix was transformed into *E. coli* DH5-α competent cells and positive clones were screened by colony PCR followed by confirmation of the cloned plasmid with BamHI/SalI and NotI/PstI restriction digestion and further transformed in BL21-CodonPlus (for pGEX-4T 1 constructs) and BL21 DE3 (for pMAL^TM^c5X constructs) *E. coli* expression strains for *Pf*PHB1 and *Pf*PHB2 recombinant protein expression.

For GST-tagged recombinant proteins, the *E. coli* cells were grown at 37 °C and were later subjected to Isopropyl β-d-1-thiogalactopyranoside (IPTG, Sigma-Aldrich, USA) induction at an OD600 ∼ 0.6-0.8 with 0.7 mM at 37 °C for 6 hours for *Pf*PHB1-pGEX-4T 1 and 0.7 mM at 16 °C for 20 hours for *Pf*PHB2-pGEX-4T 1. The *Pf*PHB1-and *Pf*PHB2-GST fusion proteins were purified via affinity chromatography from bacterial extracts using glutathione-sepharose (GE Healthcare, USA) using the soluble fractions of the lysed bacterial cultures. Whereas, for MBP-tagged recombinant proteins, the *E. coli* cultures were grown at 37 °C, subjected to IPTG induction at an OD600 ∼ 0.6-0.8 with 0.75 mM IPTG/1% Glucose and further grown for 5 hours. The bacteria cell pellets were lysed under native conditions and the recombinant *Pf*PHB1 and *Pf*PHB2 proteins were found to be expressed in the form of inclusion bodies (IB). To purify the respective recombinant proteins under denaturing conditions, the IB were harvested and further purified by standard washing conditions using detergents like Triton X-100. Successful purification of all the fusion proteins was validated by SDS-PAGE followed by Coomassie staining. Additionally, for Western blot analysis, the protein bands were transferred to Hybond ECL nitrocellulose membrane (Amersham Biosciences, UK) followed by blocking in 5% skim milk and 1% bovine serum albumin (BSA) in Tris-buffer saline (pH 7.5) followed by probing with anti-GST (dilution 1:1000) and anti-MBP (dilution 1:5000) primary antibodies in for respective recombinant proteins. After washing, the blots were probed with anti-mice secondary antibody (dilution 1:5000). The blots were developed to detect specific bands of the purified recombinant proteins. To raise specific antibodies, Balb/c mice were immunized with 100 μg of each recombinant protein in 1X phosphate buffer saline (PBS). Separate formulations were made by thoroughly mixing equal volumes of Freund’s complete adjuvant and 1X PBS containing respective recombinant proteins. For booster doses, formulations were made with Freund’s incomplete adjuvant. After the boosters were complete, respective antisera were isolated from mice blood.

### Western blot analysis for detection of protein expression

In order to detect the native protein expression pattern of *Pf*PHB1 and *Pf*PHB2 in *P. falciparum*, different asexual blood stages (rings, trophozoites, and schizonts) of the parasites at 10-12% parasitemia were harvested from synchronized cultures, while gametocyte stages (Gc) were enriched by Percoll purification. Parasites were incubated with 0.05% saponin/PBS for 10 minutes for RBCs lysis and centrifuged at 3,000 rpm for 10 minutes at 4°C followed by washing with 1X PBS. Pelleted parasites were resuspended in RIPA lysis buffer (150 mM NaCl, 0.1% Triton X-100, 0.5% sodium deoxycholate, 0.1% Sodium dodecyl sulfate (SDS), 50 mM Tris-HCl pH 8.0) supplemented with protease inhibitor cocktail (PIC, complete EDTA-free, Roche, Switzerland) and incubated for 10 minutes on ice. Lysed non-infected erythrocytes were used as a negative control. 5X SDS-PAGE loading buffer containing 25 mM dithiothreitol (DTT) was then added to the lysates, heat-denatured for 10 minutes at 95°C, and separated via SDS-PAGE followed by transferring to Hybond ECL nitrocellulose membrane (Amersham Biosciences, UK) according to the manufacturer’s protocol. Blocking of non-specific binding sites was performed by incubation with 5% skim milk and 1% BSA in Tris-buffer saline (pH 7.5) at room temperature on slow shaking. For immunodetection, membranes were incubated overnight at 4°C with polyclonal mouse anti-*Pf*PHB1 (dilution 1:500), anti-*Pf*PHB2 antisera (dilution 1:250), anti-*Pf*39 (dilution 1: 1000) antisera, in blocking solution. Following several washing steps, the membranes were incubated for 1 hour at room temperature with a goat anti-mouse alkaline phosphatase-conjugated secondary antibody (dilution 1: 5000; Sigma-Aldrich, USA) and developed in a solution of nitroblue tetrazolium chloride (NBT) and 5-bromo-4-chloro-3-indoxyl phosphate (BCIP; Roche, Switzerland) for 15 minutes at room temperature. Blots were scanned and processed using GIMP2 version 2.10.

### Immunofluorescence assays to detect protein localization

To evaluate the localization of *Pf*PHB1 and *Pf*PHB2 in *P. falciparum,* mixed asexual and gametocyte stage cultures at 8-10% parasitemia were collected and subjected to indirect immunofluorescence assays. After the cell monolayers were air-dried on glass slides, followed by fixing with 4% paraformaldehyde/PBS (pH 7.4) for 10 minutes at room temperature followed by membrane permeabilization with 0.1% Triton X-100/125 mM glycine/PBS at room temperature for 30 minutes. Blocking of non-specific binding sites was performed using 3% BSA/ PBS for 1 hour, followed by incubation with polyclonal mouse antisera against *Pf*PHB1 (dilution1:50), *Pf*PHB2 (dilution 1:50) for 1 hour at 37°C. The binding of *Pf*PHB1 and *Pf*PHB2 primary antibodies was detected by incubating the samples with monoclonal Alexa Fluor 488-conjugated goat anti-mouse IgG antibody (dilution 1:1000; Molecular Probes, USA) for 1 hour at room temperature. The different parasite stages were detected by double-labelling with stage-specific polyclonal rabbit anti-merozoite surface protein-1 (dilution 1:200; MSP-1), anti-*Pf*s230 (dilution 1:200) antisera for asexual and sexual stages, respectively, followed by incubation with monoclonal Alexa Fluor 594-conjugated goat anti-rabbit IgG antibody (dilution 1: 1000; Molecular Probes, USA). Non-immunized mice serum (dilution 1:100; NMS) was used in the assays as a negative control. Nuclei were highlighted by treatment with Hoechst33342 nuclear stain for 1 minute at room temperature, and cells were mounted with anti-fading solution AF2 (Citifluor Ltd, USA) and sealed with transparent nail paint. Immunofluorescence assays were analyzed using a Leica DM 5500B fluorescence microscope, and digital images were processed using GIMP2 version 2.10.

### Cellular fractionation of parasite lysate

By cellular fractionation of parasite cell lysates, both localization and biochemical properties of *Pf*PHB1 and *Pf*PHB2 could be assessed. For this purpose, *P. falciparum* asexual parasites with 8-10% parasitemia were harvested. After saponin-lysis and washing steps, the pellet was lysed with 1 ml lysis buffer (20 mM HEPES, pH 7.8; 10 mM KCl; 1 mM EDTA; 1 mM DTT; 1 mM PMSF; 1% Triton X-100) and incubated for 10 minutes on ice. The organellar fraction was pelleted at 5200 rpm for 5 minutes at 4°C while the supernatant constituting the cytosolic fraction was aspirated followed by washing the pellet twice with lysis buffer. For organellar protein extraction, an equal volume of extraction buffer (20 mM HEPES, pH 7.8; 800 mM KCl; 1 mM EDTA; 1 mM DTT; 1 mM PMSF; 1× PIC) was added to the pellet and incubated for 30 minutes while rotating at 4°C. The extract was cleared by centrifugation at 10,000 rpm for 30 minutes at 4°C. Supernatant containing the organellar proteins was diluted with 1 volume of dilution buffer (20 mM HEPES, pH 7.8; 1 mM EDTA; 1 mM DTT; 30% glycerol). Later, all the collected fractions were separated by SDS-PAGE and analyzed by Western blot analysis for the detection of proteins in the respective fractions. Immunoblotting was performed with primary polyclonal mouse anti-*Pf*PHB1 (dilution 1:500), mouse anti-*Pf*PHB2 (dilution 1:250), rabbit raised anti-H_4_Kac_4_ antibody (dilution 1:1000) was used as an organellar fraction positive control, while rabbit anti-NapL (dilution 1:500) served as cytoplasmic fraction control, followed by probing with respective goat anti-mouse or rabbit secondary antibodies conjugated with alkaline phosphatase (dilution 1: 5,000; Sigma-Aldrich) or conjugated with HRP (dilution 1:2000; for NapL).The blots were developed as mentioned above.

Another method was employed for subcellular fractionation, wherein parasite culture at 8-10% parasitemia was subjected to lysis in 0.03% saponin/PBS for 5 minutes at 37°C. Cells were pelleted and lysed by resuspension in 100 μl of 5 mM Tris-HCl (pH 8.0) supplemented with PIC followed by 10 minutes incubation at room temperature followed by freezing and thawing. Soluble proteins were separated by centrifugation, and the pellet was resuspended in 100 μl of 1% Triton X-100 and incubated for 30 minutes at room temperature to extract integral proteins. The final pellet constituting the insoluble proteins was resuspended in 100 μl of 0.5X PBS/4% SDS/ 0.5% Triton X-100. The samples were then subjected to Western blot as above. Immunoblotting with mouse antiserum directed against the endoplasmic reticulum-resident protein *Pf*39 (dilution 1:1000), nucleus-localized H_4_Kac_4_ and GAP45 were used as fraction controls.

### MitoTracker staining

To evaluate mitochondrial localization of *Pf*PHB proteins, asexual and sexual parasite culture was incubated with 100 nM MitoTracker Red CMXROS (M7512, Life Technologies) for 30 minutes at 37 °C, followed by washing the cells with 1X PBS. Thin smears from the parasite culture were quickly air dried and carefully fixed with 2% paraformadehyde (with 0.0075% glutaraldehyde in PBS) at room temperature for 30 minutes. The fixed cells were washed twice with 1X PBS and permeabilized using 0.1% Triton X-100 at room temperature for 3 minutes. The cells were washed twice with 1X PBS followed by blocking with 3% BSA at room temperature for 1 hour, followed by incubation with mice raised antisera against *Pf*PHB1 and *Pf*PHB2 (dilution 1:50; in 1% BSA in 1X PBS) for 1 hour at room temperature. Later, the smears were washed twice with 1X PBS-T (1X PBS in 0.05% Tween20) and once with PBS followed by incubation with monoclonal Alexa Fluor 488-conjugated goat anti-mouse IgG antibody (dilution 1:1000; Molecular Probes, USA) for 1 hour at room temperature. After washing the smear twice with PBS-T and once with PBS, nuclei were highlighted by treatment with Hoechst33342 nuclear stain for 1 minute at room temperature and processed as described above.

### Pulldown assay for *Pf*PHB1 and *Pf*PHB2 interaction

To ascertain the interaction of *Pf*PHB1 and *Pf*PHB2 with each other pulldown analysis was performed. Briefly, GST-tagged 50 µg *Pf*PHB1 or *Pf*PHB2 recombinant protein was incubated with 20 µl of GST-beads in 200 µl of binding buffer (10 mM HEPES, 150 mM NaCl, 0.5% Igepal) and incubated on rotation for overnight at 4°C for binding. Later, the sample was centrifuged at 2000 rpm for 10 minutes at 4°C, and the supernatant was saved in a fresh tube. Following washing the beads several times with 200 µl binding buffer, saponin-lysed schizont stage lysate of infected RBCs or uninfected RBC lysate was incubated with the beads for binding on rotation at 4°C overnight. The samples were centrifuged at 2000 rpm for 10 minutes at 4°C, and supernatant was transferred to fresh tubes. The beads were washed several times with binding buffer and the bound protein fractions were eluted using 20 µl of 10 mM Glutathione. The fractions were analyzed by Western blotting to decipher *Pf*PHB1 and *Pf*PHB2 as co-interacting proteins.

### MicroScale Thermophoresis (MST)

MST is a biophysical technique relying on binding-induced changes in thermophoretic mobility, depending on various molecular properties such as particle size, charge, conformation, hydration state and solvation entropy. Thereby, under constant buffer conditions, the thermophoretic mobility of unbound proteins usually differs from that of proteins bound to their interaction partners. Here, we evaluated the interaction of recombinant *Pf*PHB1 and *Pf*PHB2 proteins using Monolith NT.115 instrument (NanoTemper Technologies, Munich, Germany). Thermophoretic movement of a fluorescently labelled protein is measured by monitoring the fluorescence distribution. Shortly, 10 μM *Pf*PHB1 was prepared in 1X PBS buffer, pH 7.5, followed by labelling with 30 μM Lysine reactive dye (Monolith ^TM^ Series Protein Labelling Kit Red-NHS 2^nd^ Generation), and incubated for 30 minutes in the dark at room temperature. Following incubation, the labelled protein was passed through an equilibrated column (provided in the kit) along with the buffer. Fractions of the labelled protein were eluted and collected followed by subjecting the elutions to fluorescence count. Fluorescence counts in the range of 250-400 was taken further for interaction analysis, whilst fractions with fluorescence counts >400 were diluted as per the requirement. 10 µM *Pf*PHB2 recombinant protein, diluted in 1X PBS/0.01% Tween20, with decreasing concentrations were titrated against the constant concentration of the labelled *Pf*PHB1 protein. Samples were pre-mixed and incubated for 15 minutes, followed by centrifugation at 8000 rpm for 10 minutes at room temperature. The samples were taken into the standard treated capillaries (K002 Monolith NT.115) and thermophoretic mobility was analyzed. Similarly, to evaluate the interaction of Roc-A (Sigma-Aldrich, CAS no. 84573-16-0) with *Pf*PHB1 and *Pf*PHB2, 100 µM Roc-A, after serial dilution in 1X PBS/0.01% Tween20, was incubated with labelled *Pf*PHB1 or *Pf*PHB2 recombinant protein. Following incubation, the samples were processed as mentioned above. All experiments were carried out at room temperature, at 20% LED power and 40% MST power. Data evaluation was performed with the Monolith software (Nano Temper, Munich, Germany).

### Thermal Stability Assay (TSA)

Interaction of natively expressed *Pf*PHB1 and *Pf*PHB2 proteins with Roc-A was monitored via thermal shift assay in *P. falciparum*. Early-trophozoite stage parasites at 8-10% parasitemia were treated with 50 µM Roc-A and incubated for 4 hours at 37°C along with the untreated controls. After harvesting the parasites, cells were lysed with 0.05% saponin/PBS and the pellet was resuspended in RIPA lysis buffer. The samples were heated at different temperatures (40, 60 and 80 °C) for 6 minutes and then cooled at room temperature for 15 minutes. Following centrifugation at 10,000 rpm for 40 minutes at 4 °C, supernatant was transferred to new tubes and run on SDS-PAGE. The separated bands were transferred to a nitrocellulose membrane to probe with *Pf*PHB1 (dilution 1:500 in 3% BSA/PBS) and *Pf*PHB2 (dilution 1:250 in 3% BSA/PBS) antisera, followed by anti-mice HRP secondary antibody (dilution 1:2000) to evaluate the change in protein stability in the presence and absence of Roc-A. After developing the blots with ECL Western Blotting Substrate (Bio-rad, USA), densitometry analyses of the bands were performed with ImageJ software.

### *P. falciparum* growth inhibition assay

To assess the antimalarial activity of Roc-A and to determine half-maximal drug concentration values (IC_50_) of 3D7, R539T, RKL-9 asexual blood stage parasite inhibition, synchronized *P. falciparum* infected erythrocyte cultures were used at mid to late-ring stage (12-16 hours post-infection (h.p.i.) at a parasitemia of 0.5% and 2% hematocrit. The parasites were treated with Roc-A inhibitor (Sigma-Aldrich, St. Louis, MO, USA) for 72 hours at concentrations ranging from 0-200 nM. Untreated controls were cultured in parallel under the same conditions and processed similarly. Similarly, the NF54 strain of *P. falciparum* was used to seed gametocyte cultures at 1% parasitaemia of ring stage. Culture aliquots (approx. 0.1 to 0.2 ml) with the indicated stage of gametocytes were seeded to a 96-well plate in duplicates, and either control or test compound, Roc-A was added in different concentrations (200 nM, 100 nM, 50 nM, 10 nM, 5 nM and 0.5 nM). The culture medium was changed after 24 hours, and an appropriate concentration of compound was maintained. Blood smears of parasites were prepared after 24 and 72 hours and stained with Giemsa (Amresco, U.S.A).

To determine the inhibition in asexual and sexual stages, parasitaemia was monitored by manually counting the number of parasitized cells in estimated 2000 erythrocytes as well by SYBR Green-I (Thermo Fisher Scientific, Waltham, Massachusetts, US) fluorimetric method. To assess the total parasitemia by SYBR Green-I staining, cultures were collected after 72 hours and freeze-thawed. 100 µl lysis buffer containing 20 mM Tris (pH 7.5), 5 mM EDTA, 0.008% saponin, 0.08% Triton X-100 and 2X SYBR Green-I dye was added to the lysed parasites and incubated for 3 hours at 37°C in the dark. Fluorescence after the SYBR Green-I assay was recorded using a Varioskan Lux multi-well plate reader (Thermo Scientific) at an excitation and emission wavelength of 485 nm and 530 nm, respectively. The data were corrected for the background fluorescence of uninfected erythrocytes, normalized to the growth of control parasites. The percent inhibition was calculated with respect to untreated control. IC_50_ values of Roc-A in 3D7, R539T and RKL-9 were determined by plotting values of percent inhibition against log concentration of compound using Graphpad PRISM software.

Percent growth inhibition was calculated using the following formula:

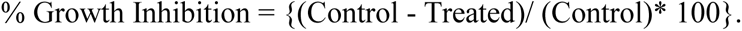

### Generation of *Pf*PHB1 and *Pf*PHB2 complementation strains in yeast mutants

Full length nucleotide sequences of *Pf*PHB1 (1-819 bp) and *Pf*PHB2 (1-915 bp) using *P. falciparum* genomic DNA template, and gene-specific primers (**Table S1**), were amplified with Phusion™ High-Fidelity DNA Polymerase (Thermo Scientific, US). The amplified DNA fragment was purified with QIAquick Gel Extraction Kit (Qiagen) using the manufacturer’s protocol. p416 GPD vector, with a host range in bacteria and yeast, harbors GPD promotor and bears Uracil-encoding gene (*ura^+^*) for selection. The purified inserts and p416 GPD vector were digested with BamHI/SalI (New England Biolabs, UK) and ligated over night at 16°C using T4 DNA ligase (New England Biolabs, UK). The ligation mix was transformed into *E. coli* DH5-α competent cells and positive clones were screened by colony PCR followed by confirmation of the cloned plasmid with BamHI/SalI restriction digestion. The *Pf*PHB1-p416 GPD and *Pf*PHB2-p416 GPD constructs were transformed in *S*. *cerevisiae* mutants SN436_Y9687::phb1Δ (YGR132C) and SN437_Y9687::phb2Δ (YGR231C), respectively using Frozen-EZ Yeast Transformation II^TM^ (Zymo Research) as per the manufacturer’s protocol and plated on YNB (yeast nitrogen base) agar plate (supplemented with 2% glucose and 1X amino acid mix without uracil) as selective media and cells were cultivated at 30°C. The transformed colonies were confirmed with colony PCR method with gene-specific primers. *S. cerevisiae* BY4742 (MATα; his3Δ; leu2Δ; lys2Δ; ura3Δ) was used as the wild type control wherever applicable.

### Growth curve analysis in complementation strains of yeast

To analyze the growth pattern of yeast mutant strains complemented with *Pf*PHB1 or *Pf*PHB2, we pre-cultured BY4742, phb1Δ and phb2Δ in YPD media (1% yeast extract, 2% peptone, 2% glucose), whilst phb1Δ-*Pf*PHB1-p416 GPD and phb2Δ- *Pf*PHB2-p416 GPD strains in YNB selective media at 30°C in the presence and absence of ethidium bromide (EtBr). EtBr was used at a concentration of 25 μg/ml, which is sufficient to destroy mitochondrial DNA of yeast cells without affecting the nuclear DNA following overnight incubation (34). Later, the pre-culture was diluted to initial A_600_ =0.05 with YPD medium and further used as the main culture. After every 20 hours, the A_600_ was measured spectrophotometrically to quantitate the growth of EtBr treated and untreated yeast cells. Additionally, following 3 days of incubation, each culture was harvested and adjusted to A_600_ = 0.5 using sterile YPD medium. The particular solution was then serially diluted 10-times and each dilution suspension was then spot on solid YPD medium and incubated for two days at 30 °C.

### Assessing effect of Roc-A on yeast mitochondrial integrity

To evaluate the effect of Roc-A on yeast cell growth and mitochondrial integrity in BY4742 wild type, phb1Δ, phb2Δ, phb1Δ-*Pf*PHB1-p416 GPD, and phb2Δ-*Pf*PHB2-p416 GPD strains were grown in their respective medium as mentioned above as pre-cultures at 30°C in the presence and/or absence of EtBr. After attaining adequate growth, the cultures were diluted to A_600_ =0.05, grown with and/or without 5 µM Roc-A at 30°C. Thereafter, growth of yeast cells was monitored at every 20 hours. Moreover, following 60 hours incubation, each culture was harvested and adjusted to A_600_ = 0.5 using sterile YPD medium. The particular solution was then serially diluted 10-times and each dilution suspension was then spot on solid YPD medium and incubated for two days at 30 °C.

## DATA AVAILABILITY

The contributions presented in the study are included in the research article/Supplementary Material, and further inquiries can be directed to the corresponding authors.

## RESULTS

### Protein architecture and transcript expression of *Pf*PHB1 and *Pf*PHB2

*In silico* analysis generated three-dimensional *Pf*PHB1 and *Pf*PHB2 protein structures predicted through I-TASSER and ModRefiner bioinformatics tools (**Figures 1A and 1B**). In order to determine the transcript expression profile of *spfh* superfamily during the blood stage life cycle, stage specific semi-quantitative reverse transcriptase PCR (RT-PCR) was performed by collecting parasites at different synchronized asexual stages, namely rings, mid to late trophozoites and late schizonts. The expression profile was also compared for gametocyte stages: immature (stages II-IV), mature (stage V) and activated (30 minutes post activation (p.a.)). The samples were subjected to RNA isolation using TRIzol reagent (Invitrogen, USA) and later used for cDNA with the SuperScript IV First-Strand Synthesis System (Invitrogen, USA) according to the manufacturer’s protocols. Transcripts of *Pfphb1* (196 bp), *Pfphb2* (155 bp), *Pfstoml* (229 bp) and *Pfphbl* (213 bp) were amplified with respective gene-specific primers (**Table S1**). High transcript levels were observed for both *Pfphb1* and *Pfphb2* in trophozoites as well as in immature and mature gametocytes stages (**Figure 1C**). However, *Pfstoml* appeared to have higher transcript expression in trophozoites as compared to the other asexual and sexual parasite stages. On the same line, *Pfphbl* also displayed lower transcript expression across all the stages with least to negligible expression in the schizonts stage. Transcript analysis for the housekeeping gene *Pfaldolase* (378 bp) encoding the enzyme aldolase was included as a loading control and showed almost equal loading of cDNA samples. The absence of gDNA in the samples was confirmed by using a control lacking reverse transcriptase. Purity of the asexual stage and gametocyte samples was demonstrated by amplifying transcripts for the asexual blood stage-specific gene *Pfama1* (189 bp) and for the gametocyte-specific gene *Pfccp2* (198 bp) (**Figure 1C**). The transcript expression of *spfh* superfamily was found to be in accordance with transcriptomic data available at the PlasmoDB database in which the transcriptome of seven stages of the malaria parasite was analyzed by RNA sequencing (PlasmoDB). The densitometry plots also revealed higher transcript expression of *Pfphb1* and *Pfphb2* in trophozoite and gametocyte stages (**Figure 1D**) as compared to others, indicating towards a possible role of these proteins in metabolically active and/or mitochondrial function-dependent stages, essential for asexual and sexual stage growth and development.

**Figure 1.**
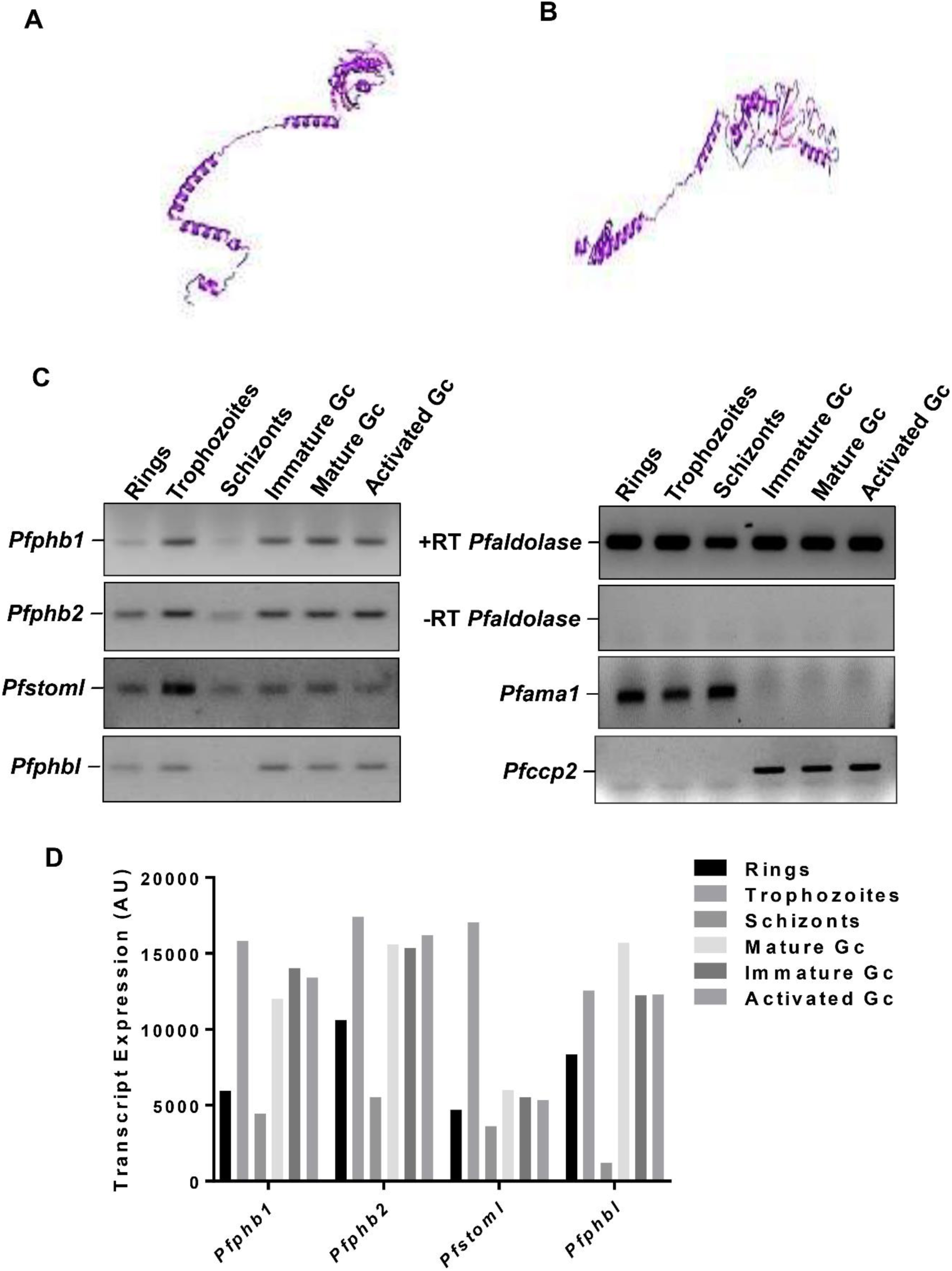
Structure prediction and transcript expression profile of *Pfphb1 and Pfphb2*. (A, B) Three-dimensional representation of predicted structures of the proteins. (C) Transcript expression of the SPFH superfamily of proteins was analysed by preparing complementary DNA (cDNA) from asexual (rings, trophozoites, schizonts) and sexual (immature, mature and 30 minutes post-activation gametocytes (Gc)) parasite stages. The cDNA was subjected to semi-quantitative reverse transcriptase (RT) PCR using gene-specific primers of *P. falciparum*. The transcript expression levels of *Pfama1* and *Pfccp2* were used to verify asexual stage and gametocyte-specific expression, respectively. cDNA samples without RT were used as controls for genomic DNA contamination, while, *Pfaldolase* with RT was used as a loading control. (D) Densitometry plots of agarose gels representing transcript expression of *Pfphb1*, *Pfphb2*, *Pfstoml*, and *Pfphbl*.

### Purification of *Pf*PHB1 and *Pf*PHB2 recombinant proteins and generation of antisera

With the aim of generating antisera and employ the purified recombinant proteins in further experiments, we cloned, expressed and purified *P. falciparum Pf*PHB1 and *Pf*PHB2 (**Figure 2A**), using affinity-based chromatography. We successfully generated full length N-terminal GST-tagged recombinant proteins of *Pf*PHB1 and *Pf*PHB2 with pGEX-4T 1 vector, whereas, with MBP-tagged recombinant proteins, specific regions of *Pf*PHB1 (115-819 bp) and *Pf*PHB2 (61-915 bp) were cloned in pMAL^TM^c5X. The purity of all the recombinant proteins was determined with by SDS-PAGE followed by Coomassie staining (**Figures 2B and 2C**). Identity of the recombinant proteins was confirmed with Western blot analysis for respective recombinant proteins with band sizes at 56.5 KDa (*Pf*PHB1-GST), 60.7 KDa (*Pf*PHB2-GST), 68 KDa (*Pf*PHB1-MBP) and 74 KDa (*Pf*PHB2-MBP) (**Figures 2D and 2E**). Polyclonal antisera were generated against these recombinant proteins in Balb/C mice.

**Figure 2.**
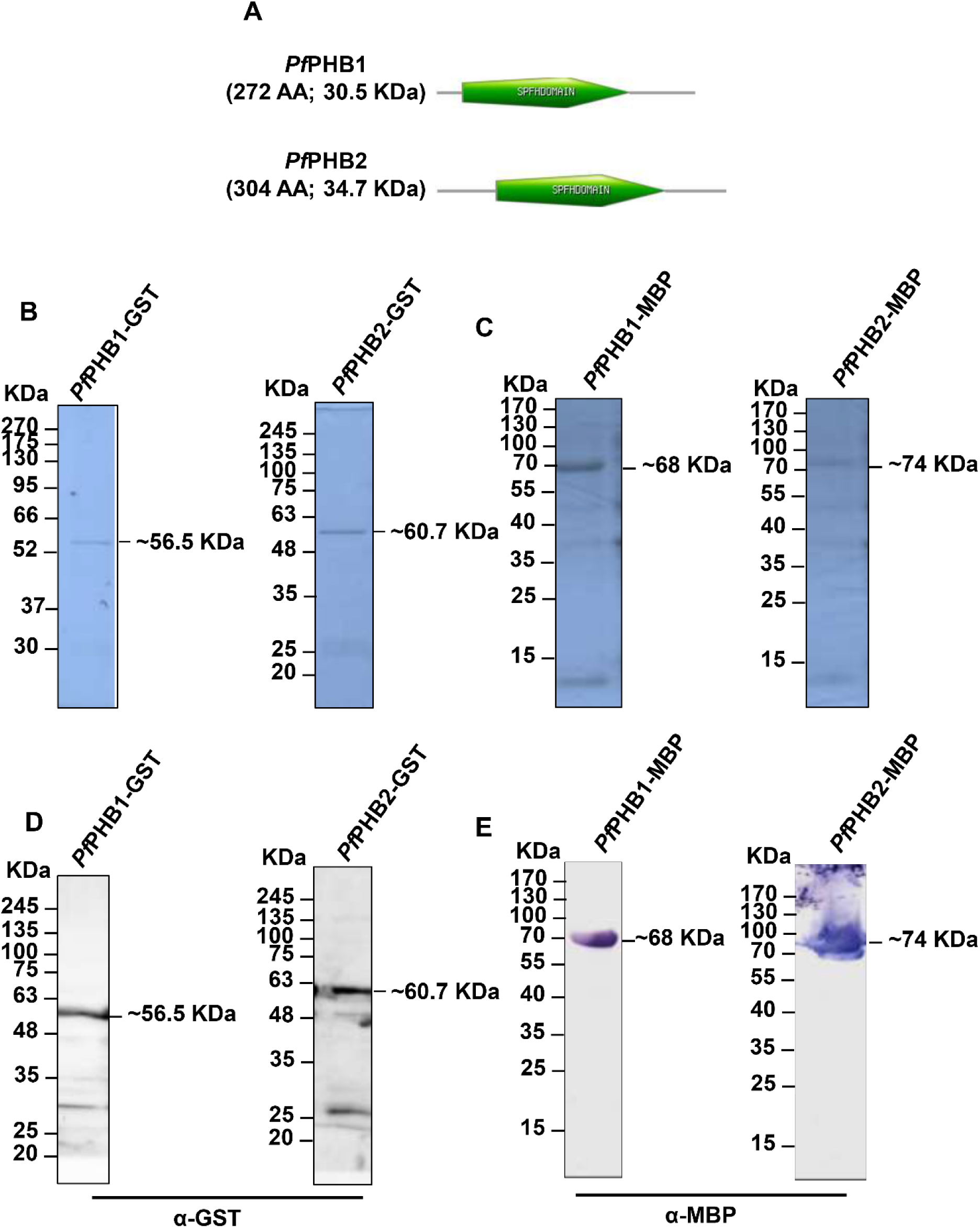
*Pf*PHB1 and *Pf*PHB2 recombinant protein purification. (A) Diagrammatic representation of *Pf*PHB1 and *Pf*PHB2, showing their SPFH (PHB/Band-7) domain (in green), expressed as recombinant proteins. (B, C) Coomassie-stained SDS-PAGE representing purified recombinant *Pf*PHB1 and *Pf*PHB2 tagged with GST (B) and MBP (C). Selected *Pf*PHB1 and *Pf*PHB2 coding regions cloned in pGEX-4T 1 and pMAL^TM^c5X vectors, after induction with appropriate concentration of Isopropyl β-d-1-thiogalactopyranoside (IPTG) were purified with affinity-binding methods. GST-tagged *Pf*PHB1 (∼56.5 KDa) and *Pf*PHB2 (∼60.7 KDa) recombinant proteins were purified with glutathione-sepharose beads. While, the MBP-tagged *Pf*PHB1 (∼68 KDa) and *Pf*PHB2 (∼74 KDa) recombinant proteins were purified from inclusion bodies using amylose-resins. (D, E) Western blot analysis of purified recombinant proteins to validate their purity and specificity. The proteins were probed with mice anti-GST (dilution 1:1000) (D) and anti-MBP (dilution 1:5000) (E) primary antibodies for respective recombinant proteins. After washing, the blots were probed with anti-mice secondary antibody (dilution 1:5000). The blots were developed to detect specific bands of the purified recombinant proteins. To raise specific antibodies, Balb/c mice were immunized with 100 μg of each recombinant protein.

### *Pf*PHB1 and *Pf*PHB2 are expressed in asexual and sexual blood stages

In an effort to decipher the expression pattern of *P. falciparum Pf*PHB1and *Pf*PHB2 proteins, we performed Western blot analysis on the parasite lysates. Synchronized *P. falciparum* parasites at 10-12% parasitemia were used to prepare cell lysates of different asexual (rings, trophozoites, and schizonts) and sexual (immature, mature, and activated gametocytes (Gc)) stages. After collecting the stage-specific lysates, the samples were subjected to SDS-PAGE followed by Western blot. The blots were probed with respective polyclonal mouse anti-*Pf*PHB1 (dilution 1:500), anti-*Pf*PHB2 (dilution 1:250) and anti-*Pf*39 (dilution 1: 1000; ∼39 KDa) antisera followed by a goat anti-mouse alkaline phosphatase-conjugated secondary antibody (dilution 1: 5,000; Sigma-Aldrich, USA) and developed in a solution of NBT and BCIP (Roche, Switzerland). The findings suggested prominent expression of *Pf*PHB1 protein in all asexual and sexual stages migrating at size of ∼30 KDa. The expression of the protein was comparable to the relative transcript levels determined for the asexual and sexual stages of *P. falciparum* with higher expression in trophozoites and gametocyte stages Incase of the *Pf*PHB2 expression, faint bands of the protein were detected in both asexual and gametocyte stages migrating at ∼34.7 KDa (**Figure 3A**). As a loading control, immunoblotting with mouse antiserum directed against the endoplasmic reticulum-resident protein *Pf*39 was used whereas, non-infected erythrocyte lysate was used as a negative control (**Figure 3A**). In conclusion, both *Pf*PHB1 and *Pf*PHB2 proteins are expressed throughout the asexual and sexual parasite stages.

**Figure 3.**
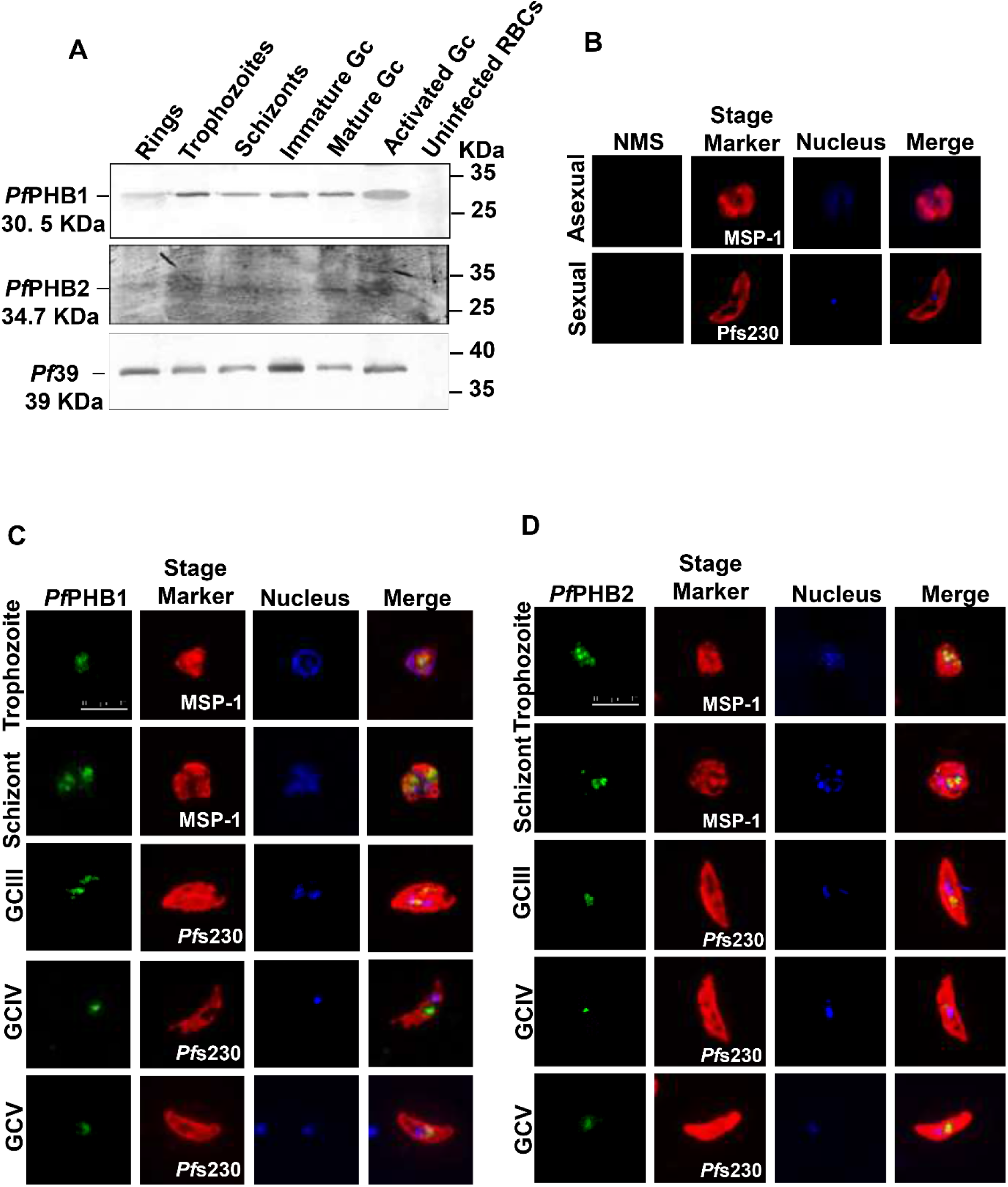
Expression of *Pf*PHB1 and *Pf*PHB2 proteins in *P. falciparum*. (A) Parasite lysates prepared from both asexual (rings, trophozoites and schizonts) and sexual (immature and mature gametocytes (Gc)) stages and gametocytes at 30 minutes p.a. (aGC) were subjected to Western blot analyses. Immunolabelling was performed with polyclonal mouse anti-*Pf*PHB1 (dilution 1:500; ∼30.5 KDa) and anti-*Pf*PHB-2 (dilution 1:250; ∼34.7 KDa) antisera to depict expression profile of the proteins. Equal loading was confirmed using a polyclonal mouse anti-*Pf*39 antiserum (dilution 1:1000; ∼39 KDa). Lysate of uninfected erythrocytes was used as a negative control. (B) Immunofluorescence assay was performed as a negative control on asexual (schizonts) and sexual (Stage V) stages. Parasites were incubated with non-immunized mice sera (NMS; dilution 1:100; green) and counter labelled with rabbit antibodies directed against-merozoite surface protein-1 (*Pf*MSP-1; dilution 1:200) and *Pf*s230 (dilution 1:200) as indicated (red). Nuclei were highlighted with Hoechst33342 nuclear stain (blue). (C, D) Immunofluoroscence assays to decipher expression and sub-cellular localization of *Pf*PHB1 (C) and *Pf*PHB2 (D) in asexual (trophozoites and schizonts) and sexual (stage III-V) stages of the parasite via immunolabelling with polyclonal mouse anti-*Pf*PHB1 (dilution 1:50) and anti-*Pf*PHB2 (dilution 1:50) antisera (green). Asexual stages were highlighted with polyclonal rabbit anti-*Pf*MSP-1 (dilution 1:200) antibody, whereas, gametocyte stages were marked with rabbit anti-*Pf*s230 (dilution 1:200) antibody (red). Nuclei were marked with Hoechst33342 nuclear stain (blue). Bar, 10 μm. Results are representative of three independent experiments.

To further validate the expression pattern of PHBs in asexual as well as sexual blood stages of *P. falciparum*, we performed indirect immunofluorescence assays. Asexual (trophozoites, and schizonts) and sexual (gametocyte stage III-V) stage parasite cultures at parasitemia of 8-10% were immunolabelled with respective mouse anti-*Pf*PHB1 and anti-*Pf*PHB2 antisera followed by Alexa-Fluor 488 conjugated goat anti-mouse IgG antibody (Invitrogen, USA). It demonstrated expression of *Pf*PHB1 and *Pf*PHB2 proteins in trophozoites and schizonts as well as in immature (gametocyte stage III and IV) and mature gametocytes (gametocyte stage V) of *P. falciparum*. The expression of these proteins was focused as single dots close to the nucleus, which might represent the single mitochondrion of the parasite in the blood stages, with almost no signal in other cellular compartments (**Figures 3C and 3D**). Polyclonal rabbit antisera targeted against MSP-1 and *Pf*s230 were used as stage markers for asexual and sexual stages, respectively, followed by probing with Alexa-Fluor 594-conjugated goat anti-rabbit IgG antibody (Invitrogen, USA) (**Figures 3C and 3D**). When non-immunized mice serum (NMS; dilution 1:100) was used in the assays as a negative control, no labelling was detectable in either asexual or gametocyte stages (**Figure 3B**). These data together largely validate the expression profile throughout the asexual and sexual stages of the parasite along with proposing their localization in the parasite mitochondria.

### *Pf*PHB1 and *Pf*PHB2 are organellar proteins with mitochondrial localization

PHB1 and PHB2 have been identified to be integral membrane proteins of the mitochondrial inner membrane mainly in eukaryotic cells (9–11). However, location of these proteins in *P. falciparum* is yet to be elucidated. In line with this, one such study on murine malaria model parasite *Plasmodium berghei* proposed localization of *Pb*PHB1 and *Pb*PHB2 essential proteins in mitochondria (35). To determine the nature of localization of PHB proteins, we performed sub-cellular fractionation. Asexual trophozoite lysate of the parasite was subjected for extraction of organellar and cytosolic fractions. The extracted protein fractions were then separated on SDS-PAGE followed by Western blot analysis. Immunoblotting was performed with primary polyclonal mouse anti-*Pf*PHB1 (dilution 1:500), mouse anti-*Pf*PHB2 (dilution 1:250), mouse anti-NapL (dilution 1: 500), rabbit anti-H_4_Kac_4_ antibody (dilution 1: 1000) followed by probing with respective goat anti-mouse or rabbit secondary antibodies conjugated with alkaline phosphatase (dilution 1: 5000) or HRP (dilution 1:2000). After developing the blots, it suggested that *P. falciparum Pf*PHB1 (∼30 KDa) and *Pf*PHB2 (∼34.7 KDa) are organellar protein with almost no expression in the cytosolic fraction (**Figures 4A and 4B**). This observation was found to be in accordance with the immunolabelling where *Pf*PHBs were found to be expressed and localized close to the nucleus of the parasite, with a possible mitochondrial origin. Nuclear protein H_4_Kac_4_ (∼11 and ∼13 KDa) and cytosolic protein NapL (∼ 40 KDa) were used as markers for cellular fractionation of proteins. Additionally, another approach of subcellular fractionation was employed to extract soluble, peripheral, integral membrane and insoluble protein fractions. Interestingly, the *Pf*PHB1 showed expression prominently in the integral membrane fraction at size ∼30 KDa. Immunoblotting with mouse antiserum directed against the endoplasmic reticulum-resident protein *Pf*39, nucleus-localized H_4_Kac_4_ and integral membrane protein GAP45 were used as fraction controls (**Figures 4C**). The blots were scanned and processed using Adobe Photoshop CS5 software.

**Figure 4.**
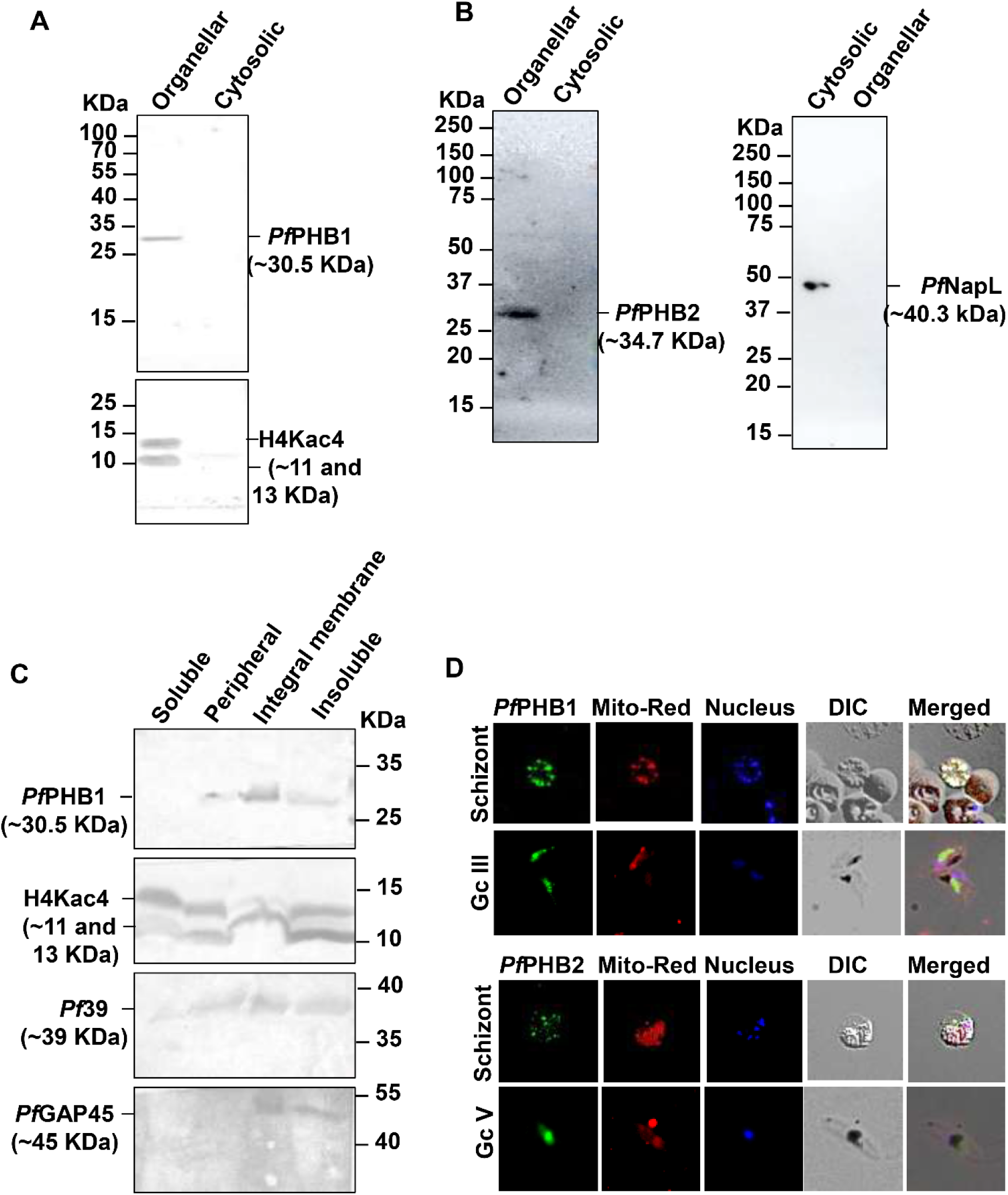
*Pf*PHBs are organellar proteins with mitochondrial localization. (A) Sub-cellular localization of *Pf*PHB1 and *Pf*PHB2 were detected in cytosolic and organellar fractions of enriched asexual parasites (trophozoites) which were subjected to Western blot analysis probed with their respective mice antisera. Bands were detected at ∼30.5 KDa (A) and ∼34.7 KDa (B) for *Pf*PHB1 and *Pf*PHB2, respectively, in organellar fractions. Rabbit antibody against H4Kac4 (dilution 1:1000) detecting acetylated histone H4 (∼11 and ∼13 KDa) was used as organellar fraction control while rabbit anti-NapL antisera (1:500) with size at ∼40.3 KDa. (C) Asexual stage (trophozoite) lysate of the parasite was used to extract soluble, peripheral, integral membrane and insoluble protein fractions. The samples were subjected to Western blot and immunolabelled with mouse anti-*Pf*PHB1 antisera to detect *Pf*PHB-1 (∼30.5 KDa). Rabbit anti-H4Kac4 (dilution 1:1000) antisera (∼11 and ∼13 KDa), mouse anti-*Pf*39 (dilution 1:1000) antisera (∼39 KDa) and rabbit anti-*Pf*GAP45 (dilution 1:100) antisera (∼45 KDa).

To further validate the mitochondrial localization of *Pf*PHB1 and *Pf*PHB2, we co-localized the proteins with MitoTracker Red CMXRos dye in the asexual and sexual stages of the parasite. It is a red-fluorescent dye that stains mitochondria in live cells and its accumulation is dependent upon the membrane potential. The dye is well-retained after aldehyde fixation. The mitochondria displayed efficient labelling with MitoTracker staining, largely confining to the organelle, along with co-localization with *Pf*PHB1 and *Pf*PHB2 proteins (**Figure 4D)**. Altogether, cellular fractionation and MitoTracker staining supports *Pf*PHB1 and *Pf*PHB2 as mitochondrial-resident proteins.

### *Pf*PHB1 and *Pf*PHB2 interact with each other

The interaction of PHB1 and PHB2 has been reported in previous studies where these proteins via the C-terminal coiled-coil domain mediates the interaction (9). In order to validate this interaction, we performed pulldown assay and MST analysis. *Pf*PHB1 and *Pf*PHB2 were detected by pulling down with purified recombinant *Pf*PHB2 and *Pf*PHB1 proteins from *P. falciparum* cell lysate, respectively. Briefly, GST-tagged recombinant *Pf*PHB1 or *Pf*PHB2 proteins (50 µg) were bound with GST-beads followed by washing of the beads with PBS to remove unbound proteins. Parasite lysate was incubated with the beads at 4°C overnight and the beads were further washed to remove unbound proteins. Binding of *Pf*PHB2 or *Pf*PHB1 was detected through Western blotting using anti-mouse polyclonal antibodies raised against the proteins. Bands of native *Pf*PHB1 and *Pf*PHB2 were observed when pulled down with recombinant *Pf*PHB2 and *Pf*PHB1, respectively, subsequently probing with respective anti-mouse antibodies. GST-bead bound *Pf*PHB1 or *Pf*PHB2 incubated with uninfected RBCs were used as control (**Figures 5A and 5B**).

**Figure 5.**
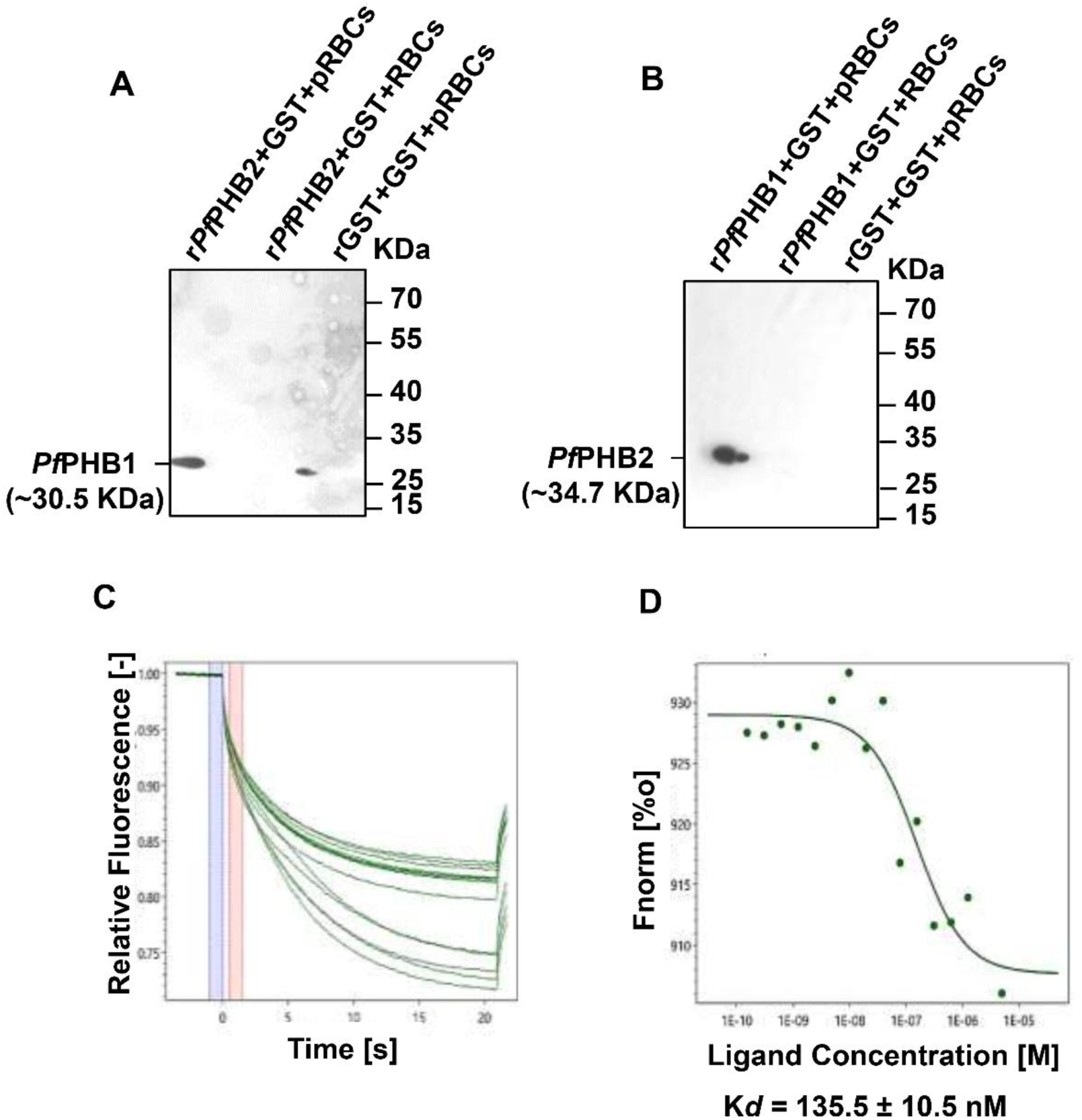
*Pf*PHB1 and *Pf*PHB2 show interaction with each other. (A, B) Western blot analysis representing pulling down of *Pf*PHB1 (A) and *Pf*PHB2 (B) from parasite lysate using recombinant *Pf*PHB2 and *Pf*PHB1 proteins, respectively. Briefly, the recombinant proteins were incubated with GST-beads in binding buffer followed by incubation with the parasite lysate (pRBCs). Incubation with uninfected RBCs (RBCs) was used as a control. The unbound proteins were removed through centrifugation and the proteins bound to the GST-beads were eluted using 10 mM Glutathione. The blots were probed with mice antisera against *Pf*PHB1 (dilution 1:500; ∼30.5 KDa) and *Pf*PHB2 (dilution 1:250; ∼34.7 KDa) followed by rabbit anti-mice HRP-conjugated secondary antibody (dilution 1:2000). (C, D) Dissociation (C) and dose-response (D) curves representing interaction analysis of *Pf*PHB1 with *Pf*PHB2 using MST. 10 µM *Pf*PHB2 recombinant protein, diluted in 1X PBS/0.01% Tween20, with decreasing concentrations were titrated against the constant concentration of the labelled *Pf*PHB1 recombinant protein. K*d* value was determined to be 135.5 ± 10.5 nM for interaction of *Pf*PHB1 with *Pf*PHB2.

Studying protein-protein interactions by biophysical approaches like MST will be another reliable technique to establish bio-molecular interactions, by exploiting immobilization-free analysis to assess the interaction of fluorescently-labelled proteins with its probable ligands/proteins. In order to accomplish this, fluorescently labelled GST-tagged recombinant *Pf*PHB1 was titrated against decreasing concentrations of GST-tagged *Pf*PHB2, where thermophoretic mobility significantly changed supporting effective interaction between the two proteins. Dissociation constant, K*d*, was found to be 135.5 ± 10.5 nM for this interaction suggesting a strong affinity between the two entities (**Figures 5C and 5D**). Together, the above findings support the possible interaction of *Pf*PHB1 and *Pf*PHB2 forming a heterodimeric structure at the inner mitochondrial membrane of the parasite.

### Roc-A inhibits *P. falciparum* growth by targeting *Pf*PHB1 and *Pf*PHB2

Roc-A has been identified as a putative interacting partner of PHB proteins in cancer cells by binding to either PHB1 and/or PHB2 (23). However, its interaction with *Pf*PHB1 and *Pf*PHB2 has not been elucidated yet. So, in order to establish these interactions, we performed both *in-vitro* and *in-vivo* assays through microscale thermophoresis (MST) and thermal stability assay (TSA), respectively. We performed MST analysis with recombinant proteins and Roc-A as their ligand. It is a novel technique to analyze interactions in an immobilization-free environment *in vitro*. When fluorescently labelled recombinant *Pf*PHB1 and *Pf*PHB2 were titrated against decreasing concentrations Roc-A, their thermophoretic mobility significantly changed suggesting effective interactions with the compound. Much to our interest, K*d* values of 1.37 ± 0.6 µM and 0.683 ± 0.021 µM were observed for interaction of *Pf*PHB1 and *Pf*PHB2 with Roc-A, respectively (**Figures 6A-6D**). These low K*d* values indicate strong affinity of interaction between *Pf*PHB proteins towards Roc-A compound.

**Figure 6.**
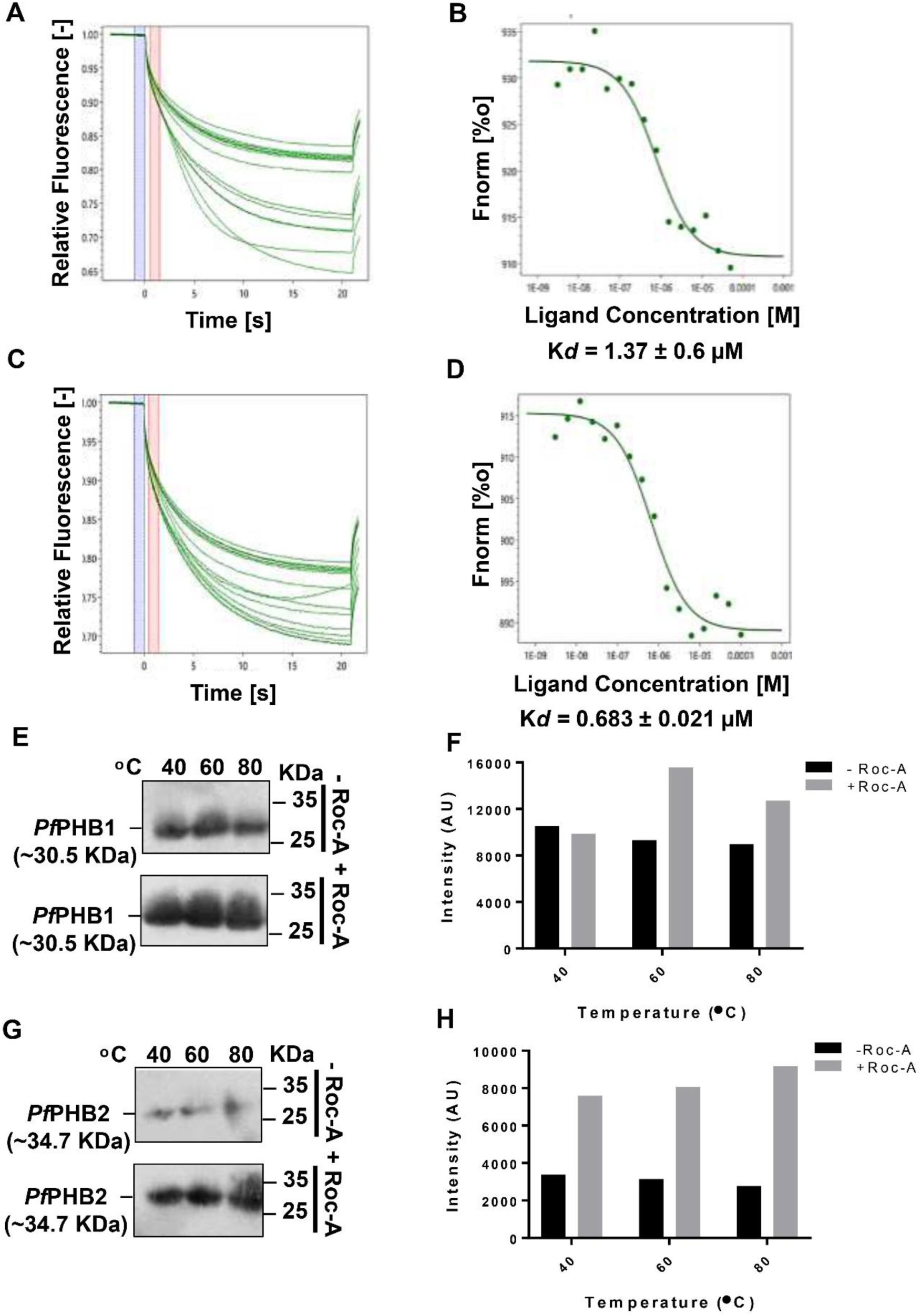
Interaction of Rocaglamide (Roc-A) with *Pf*PHB1 and *Pf*PHB2. (A-D) Dissociation (A,C) and dose-response (B, D) curves representing interaction analysis of recombinant proteins *Pf*PHB1 (A,B) and *Pf*PHB2 (C,D) with Roc-A through MST. Labelled *Pf*PHB1 and *Pf*PHB2 recombinant proteins, diluted in 1X PBS/0.01% Tween20, were titrated against decreasing concentrations 100 µM Roc-A. K*d* values were determined to be 1.37 ± 0.6 µM and 0.683 ± 0.021 µM for interaction of *Pf*PHB1 and *Pf*PHB2, respectively. (E,G) Western blots representing thermo stability analysis to validate interaction of Roc-A with native *Pf*PHB1 and *Pf*PHB2. To serve this purpose, trophozoite stage parasites were treated with 50 µM Roc-A followed by incubation for 4 hours along with the untreated control. After harvesting the parasites, cells were lysed and the pellets were resuspended in RIPA lysis buffer followed by heating the samples at different temperatures (40, 60 and 80 °C) for 6 minutes. Following centrifugation at 10,000 rpm for 40 minutes at 4°C, supernatant was transferred to new tubes and were subjected to Western blot to evaluate the change in protein stability in the presence and absence of Roc-A. (F,H) Histogram plot representing densitometry analysis of the bands performed with ImageJ software for the TSA of *Pf*PHB1 and *Pf*PHB2 with Roc-A.

Next, to further validated the above findings, thermostability of proteins was determined by TSA, relying on the fact that protein-ligand interaction bestows significant changes in the thermodynamic features of the protein leading to altering its stability with simultaneous increase in temperature. Interaction of native *Pf*PHB1 and *Pf*PHB2 with Roc-A (**Figures 6E and 6G**) was elucidated through TSA at different temperatures ranging from 40°C to 80°C. Strikingly, *P. falciparum* trophozoite stage saponin-lysed parasite lysate treated with 50 µM Roc-A showed higher protein abundance of *Pf*PHB1 and *Pf*PHB2 at higher temperature as compared to its DMSO treated counterparts when probed with respective antisera. These findings demonstrate notable thermo stabilization of the proteins in ligand-bound condition in presence of Roc-A. Densitometry plots also described similar patterns (**Figures 6F and 6H**). Taken together, these results propose Roc-A as a potent interacting ligand of PHB proteins in *P. falciparum*.

### Roc-A significantly inhibits parasite growth

After establishing the interaction of PHBs with Roc-A in *P. falciparum*, we were interested to decipher the antimalarial properties of the compound. To do so, growth inhibitory activity of Roc-A was evaluated *in-vitro* in *P. falciparum* 3D7 (chloroquine-sensitive), R539T (artemisinin-resistant) and RKL-9 (chloroquine-resistant) strains. 3D7, R539T and RKL-9 cultures at 0.5% parasitemia with highly synchronized parasites at ring stage (12-16 hours post infection) were treated with different concentrations of Roc-A (1-200 nM) in complete RPMI medium for the next 72 hours to assess parasite growth inhibition. The parasite stages were monitored by analyzing Giemsa-stained smears. It was observed that Roc-A was effectively inhibiting parasite growth in nanomolar ranges in 3D7 (IC_50_ 1.579 ± 0.22 nM), R539T (IC_50_ 4.87 ± 1 nM) and RKL-9 (IC_50_ 78.88 ± 34 nM) strains (**Figures 7A-7D**). Moreover, the inhibitory effect of Roc-A resulted into observable and significant morphological changes in the parasite by forming ’pyknotic’ bodies in the treated cultures (**Figures 7A-7C**). Additionally, the antimalarial activity of Roc-A was also elucidated in gametocyte stages of the parasite. To achieve this, gametocytes were treated with Roc-A (0.5-200 nM). Upon 24 hours incubation with compound at a lowest 10 nM concentration, majority of the different stages of gametocyte found to be morphologically distorted and abnormal compared to the untreated control (**Figure 7E)**. Furthermore, majority of abnormal stage II gametocytes did not progress to stage-III and V, respectively, therefore percent gametocytemia of mature stages (III & V) was found to be lower throughout the time period Following 24 hours treatment, gametocytemia reduced to < 0.1% in the treated parasites compared to control where gametocytemia reached 1.2%. Furthermore, with 72 hours treatment with Roc-A, majority of the gametocytes attained abnormal morphology and were unable to progress further in the cycle (**Figure 7F**). These data cumulatively demonstrate that Roc-A was effective to control the gametocytes stage II-V progression as well as reduced the gametocyte percentage *in vitro.* Thus, we propose Roc-A as a potent naturally occurring compound with antimalarial properties targeting malaria parasite by inhibiting *Pf*PHB1 and *Pf*PHB2, which eventually results in parasite growth inhibition.

**Figure 7.**
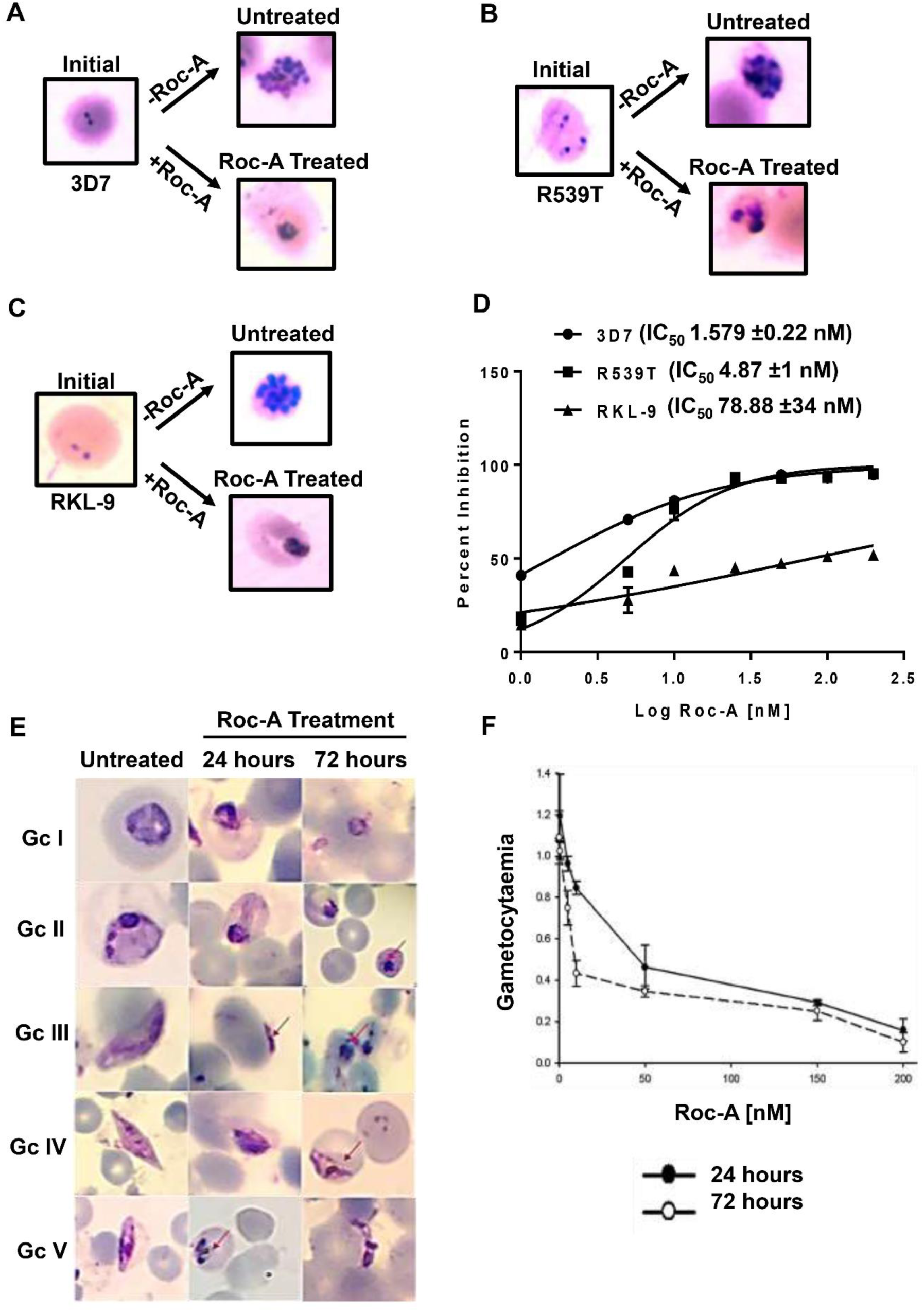
Roc-A as a potent antimalarial agent. (A-C) Giemsa stained images of initial cultures of *P. falciparum* 3D7 (A), R539T (B) and RKL-9 (C) treated with Roc-A for 72 hours forming ‘pyknotic bodies’. Cultures without Roc-A treatment were considered as controls. representing percent inhibition caused by antimalarial activity of Roc-A at a variable concentration scale. 3D7, R539T and RKL-9 cultures at 0.5% parasitemia ring stage were treated with different concentrations of Roc-A (1-200 nM) in complete RPMI medium for 72 hours to assess parasite growth inhibition. (D) Roc-A exhibits IC_50_ values in nanomolar ranges *in-vitro* in *P. falciparum* 3D7 (chloroquine-sensitive; IC_50_ 1.579 nM±0.22 nM), R539T (artemisinin-resistant; IC_50_ 4.87 nM±1 nM) and RKL-9 (chloroquine-resistant; IC_50_ 78.88 nM±34 nM) strains as represented through the log graph. (E) Giemsa-stained images of different stages of gametocytes (Gc I-V) treated with Roc-A for 24 and 72 hours. Gametocytes without Roc-A treatment were used as a control. (F) Line graph highlighting gametocidal activity of Roc-A. Gametocytes were treated with Roc-A at a concentration range (0.5-200 nM) for 24 and 72 hours.

### *Pf*PHB complementation in yeast mutants rescues cell growth

As we elucidate the mitochondrial role of *Pf*PHB in *P. falciparum*, we exploited an orthologous system, *S. cerevisiae*, to prove our hypothesis as the dispensable function of mitochondria makes the organism vulnerable to certain conditions altering the homeostasis of the organelle. To explore the *Pf*PHBs functional role in stabilizing the mitochondria and maintaining the mitochondrial integrity, an ethidium bromide (EtBr) based study was carried out. EtBr is known to selectively degrade mitochondrial DNA without disrupting the genomic DNA of yeast cells (34). When phb1Δ and phb2Δ mutants of yeast were complemented with *Pf*PHB1-p416 GPD and *Pf*PHB2-p416 GPD, respectively, the growth of the complemented strains was restored as compared to the phb1Δ and phb2Δ mutant strains (**Figure 8A**). Moreover, even in the presence of EtBr, yeast mutant strains complemented with *Pf*PHBs displayed more growth as compared to the non-complemented cells as reflected through the growth curve (**Figure 8B**). Similarly, spot assay findings corroborated with the growth curve where the complementation of *Pf*PHB1 and *Pf*PHB2 in yeast mutants displayed enhanced growth in the presence as well as in the absence of EtBr (**Figures 8C and 8D**). In gist, the effect of EtBr was more prominent in mutants as compared to the wild type indicating the absence of yeast PHBs resulted in loss of mtDNA thereby reducing the growth of yeast cells. However, this effect was successfully reversed when *Pf*PHBs were expressed in mutants restoring the loss of mtDNA leading to enhanced growth. Thus, we can infer that *Pf*PHBs have significant role in maintaining the integrity of mtDNA.

**Figure 8.**
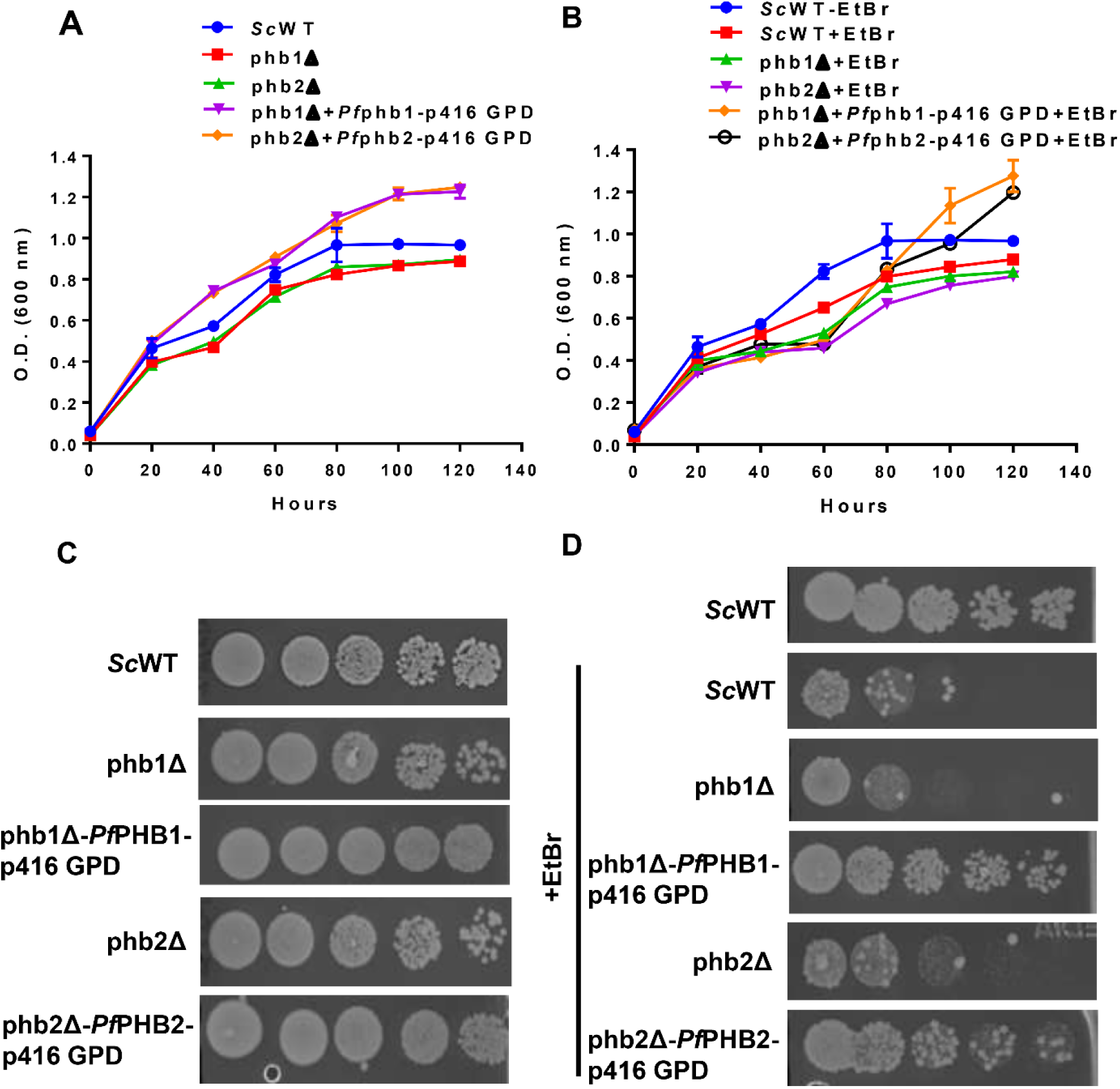
Rescue of cell growth through yeast complementation of *Pf*PHB1 and *Pf*PHB2. (A, B) Line graph reflecting growth curve of *S. cerevisiae* BY4742 wild type (*Sc*WT), phb1Δ and phb2Δ yeast mutants, and yeast complemented phb1Δ-*Pf*phb1-p416 GPD, phb2Δ-*Pf*phb2-p416 GPD strains in the absence (A) and presence (B) of EtBr. The yeast cells were pre-cultured with/without EtBr and later diluted to initial A_600_ =0.05 with YPD medium to be used further as the main culture. After every 20 hours, the A_600_ was measured spectrophotometrically to quantitate the growth of EtBr treated and untreated yeast cells. (C, D) Spot assay showing growth of yeast cells spotted following 3 days of incubation. Each culture was harvested and adjusted to A_600_ = 0.5 using sterile YPD medium. The particular solution was then serially diluted 10-times and each dilution suspension was then spot on solid YPD medium and incubated for two days at 30 °C.

### Roc-A targets *Pf*PHBs in yeast complementation

As we highlighted the significance *Pf*PHB complementation in rescuing yeast cell growth in EtBr treated cells of the mutants, the next question to be answered was whether Roc-A could target *Pf*PHB in yeast complementation model. To achieve this aim, BY4742 wild type, phb1Δ, and phb2Δ mutants and *Pf*PHB1-p416 GPD, and *Pf*PHB2-p416 GPD complemented yeast strains were grown in the presence and/or absence of EtBr and/or Roc-A, to evaluate the growth patterns. Interestingly, yeast cells of wild type, phb1Δ, phb2Δ, phb1Δ-*Pf*PHB1-p416 GPD, and phb2Δ-*Pf*PHB2-p416 GPD complemented yeast strains, in the presence of 5 µM Roc-A displayed reduced growth as compared to the EtBr and Roc-A untreated control (**Figures 9A and 9B**). Much to our surprise, in case of phb1Δ-*Pf*PHB1-p416 GPD, and phb2Δ-*Pf*PHB2-p416 GPD complemented yeast strains, the cells displayed reduced growth in the presence of Roc-A which could be primarily attributed to the drug-targeting effect of Roc-A towards *Pf*PHBs. Similarly, in the spot assay analysis, the phb1Δ-*Pf*PHB1-p416 GPD, and phb2Δ-*Pf*PHB2-p416 GPD complemented yeast strains treated with Roc-A did not form as many colonies as compared to the untreated control (**Figures 9C and 9D**). Taken together, these observations lead to affirming the fact that Roc-A targets *Pf*PHBs and causes growth defects in the cells which could be possibly due to disruption of the mitochondrial integrity and homeostasis.

**Figure 9.**
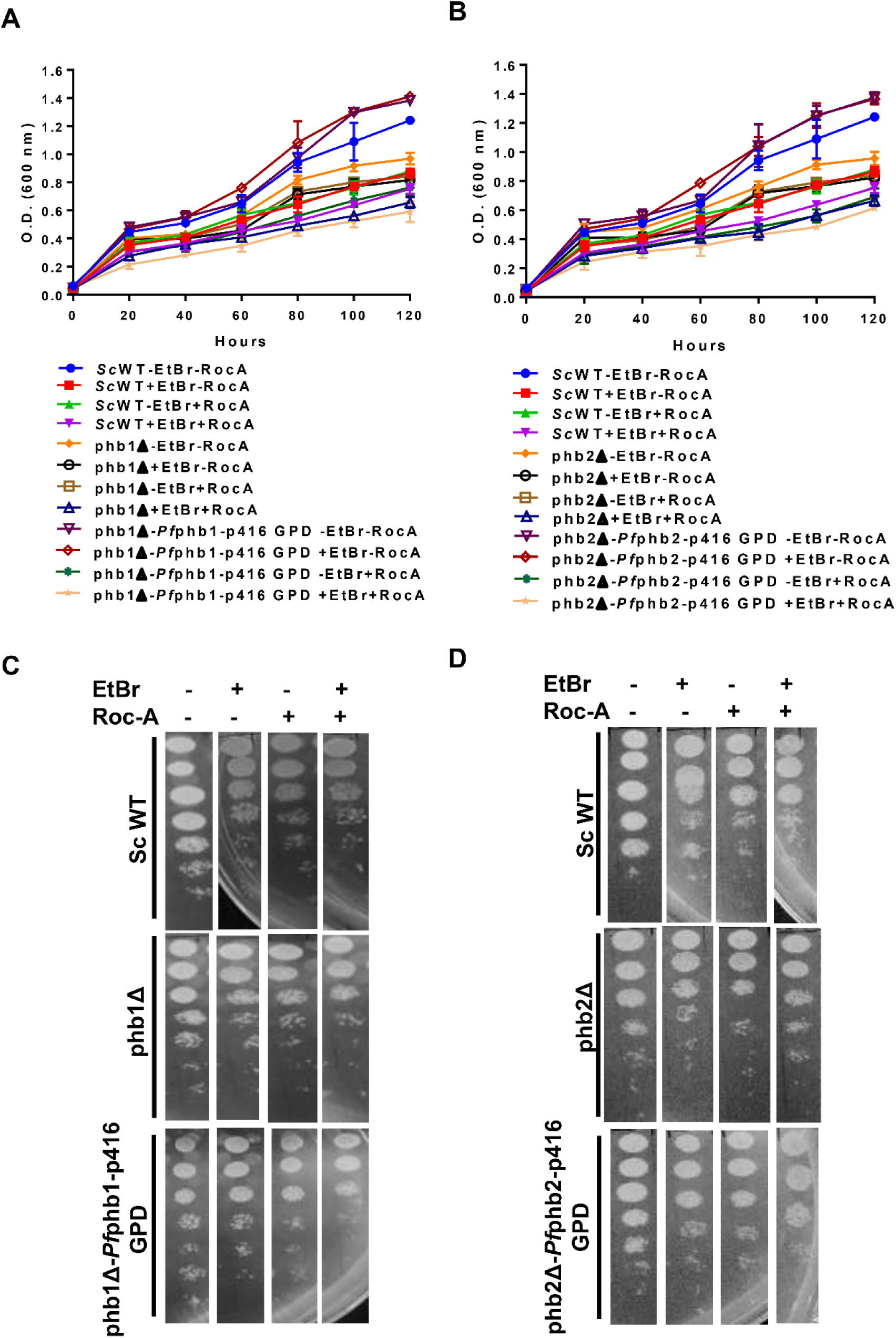
Drug targeting effect of Roc-A towards *Pf*PHBs. (A, B) Line graph showcasing growth of yeast cells of *S. cerevisiae* BY4742 wild type (*Sc*WT), phb1Δ and phb2Δ yeast mutants, and yeast complemented phb1Δ-*Pf*phb1-p416 GPD (A), phb2Δ-*Pf*phb2-p416 GPD (B) strains in the absence and/or presence of EtBr and/or Roc-A. Roc-A and EtBr were used at a concentration of 6 µM and 25 µg/ml, respectively. The yeast cells were pre-cultured with/without EtBr and later diluted to initial A_600_ =0.05 with YPD medium to be used further as the main culture in the presence and absence of Roc-A. After every 20 hours, the A_600_ was measured spectrophotometrically to quantitate the growth of yeast cells. (C, D) Spot assay showing growth of yeast cells spotted following 2 days of incubation. Each culture was harvested and adjusted to A_600_ = 0.5 using sterile YPD medium. The particular solution was then serially diluted 10-times and each dilution suspension was then spot on solid YPD medium and incubated for two days at 30 °C.

## DISCUSSION

The role played by SPFH superfamily of proteins in various cellular and mitochondrial functions has been vastly elaborated and appreciated. PHB1 and PHB2, two essential members of this protein family, are highly conserved and form a heterodimeric ring-like structure in the inner mitochondrial membrane (10, 11). PHBs display significant roles assembly and activity of the oxidative phosphorylation system (OXPHOS), mitochondrial biogenesis, mitochondrial apoptosis, mitophagy, and mitochondrial respiratory chain subunit degradation along with maintaining the overall mitochondrial homeostasis (13–16, 36). Hence, due to multiple functions in mitochondria, PHBs have been found to be as potential targets for therapeutic approaches. Though the PHBs are vastly studied in higher eukaryotic systems, their characterization and significance in *P. falciparum* is highly underappreciated.

Interestingly, PHB proteins have higher expression in cells with predominant mitochondrial functions departing more susceptibility towards mitochondrial dysfunction. In *P. falciparum*, structural and functional variations in the mitochondria of asexual and sexual (gametocytes) blood stages exits. The *Plasmodium* species harbor a single mitochondrion, that gets developed into tubular mitochondrial cristae in the gametocytes which are not present in the asexual stages but are required for complete mitochondrial functions. All these facts indicate that gametocytes might prominently rely on mitochondria for their energy requirements have a higher demand for energy transduction making them more metabolically active than the asexual stages (37). With a more active mitochondria generating higher energy in the gametocyte stage may be significant for their survival during transmission into the mosquito vector. While in the asexual stage, the parasite relies exclusively on cytosolic glycolysis and abstain from the mitochondrial tricarboxylic acid (TCA) cycle (38). With this background knowledge, we have made an attempt to study the significance of *Plasmodium* mitochondria in a bit more depth through characterizing *Pf*PHB proteins. Despite the fact that the *Plasmodium* genome encodes PHBs, their role is still unclear in the cell biology and/or signaling pathways of the parasite.

Most of mitochondrial proteins after getting transcribed and translated in the nucleus, are imported into the organelle, with an unclear understanding of the mitochondrial proteome of *Plasmodium*. As *Pf*PHBs are encoded by the nuclear DNA of the parasite, the transcript expression was found to be consistent throughout the blood stages with higher expression in trophozoite and gametocyte stages implying their essentiality in these stages (**Figures 1C and 1D**). In line with this, protein expression pattern of *Pf*PHB1 and *Pf*PHB2 was also found to be quite similar as the transcript expression, with a slightly higher expression in the trophozoite and gametocyte stages (**Figure 3A**). Additionally, the expression of these proteins was focused as single dots close to the nucleus, which might represent the single mitochondrion of the parasite in the blood stages, with almost no signal in other cellular compartments (**Figures 3C and 3D**). These data, in conjunction, indicate towards the significance of the functional aspect of *Pf*PHB proteins in these stages.

With a potential localization in the *Plasmodium* mitochondria (35), we assessed *Pf*PHBs through cellular fractionation and MitoTracker staining to ascertain the same. Much to our expectation, both *Pf*PHB1 and *Pf*PHB2 were expressed in the organellar fraction with almost no expression in the cytosolic fraction (**Figures 4A and 4B**). Additionally, *Pf*PHB1 was found to be expressed in the integral membrane fraction as well (**Figure 4C**). Validating their mitochondrial localization was the next question in line which was answered via staining the parasites with MitoTracker dye which co-localized with *Pf*PHB1 and *Pf*PHB2 expressing in the mitochondria of the parasite (**Figure 4D)**. A recent study in *P. berghei* revealed co-localization with the mito-GFP marker of PHB2, STOML, and PHBL localizing to the parasite mitochondrion (35), corroborating with our findings.

Revealing the location of both *Pf*PHBs to the *Plasmodium* mitochondria indicated towards the possibility of these proteins to interact with each other as many studies in the cancerous and other eukaryotic cells have suggested similar findings where PHB1 and PHB2 interact through their C-terminal coiled-coil domain to form a complex in the inner mitochondrial membrane (9). In line with this, we confirmed prominent interaction of *Pf*PHB1 and *Pf*PHB2 through pull down methods and MST (**Figure 5**) with the probability of forming a functional complex at the inner mitochondrial membrane. These findings strongly propose *Pf*PHBs to serve as a drug target in the mitochondria for the anti malarials. Interestingly, some antimalarial drugs are known to target specifically the mitochondrion, including, atovaquone (39–42), primaquine (43, 44), tetracycline (45), and artemisinin derivatives (46, 47), supporting these drugs as mitochondrion-targeting chemotherapeutics. We propose Roc-A as a potent ligand of *Pf*PHB1 and *Pf*PHB2 (**Figure 6)** which could target the mitochondria of the parasite. Antimalarial activity of Roc-A was evaluated in different strains of *P. falciparum* with IC_50_ values in the nanomolar ranges (**Figure 7**). Additionally, inhibition of gametocytes by Roc-A led to morphological aberrations which could be imparted through mitochondrial membrane potential disruption as an effect of inhibiting *Pf*PHB1 and *Pf*PHB2 (35). Thus, characterizing *Pf*PHBs could add to the already available information regarding the mitochondria proteome of *Plasmodium*.

Apart from studying the *Pf*PHBs in the *Plasmodium*, orthologous *S. cerevisiae* system was exploited to serve our purpose. The ability of *Pf*PHBs to prevent mitochondrial DNA damage, in yeast mutants after complementation, in the presence of EtBr clearly indicated their role in maintaining the mitochondrial integrity of the yeast cells and proposed their potential as a drug target. The growth curve reflected a higher growth pattern for mutants expressing *Pf*PHB1 and *Pf*PHB2 in the absence as well as presence of EtBr (**Figure 8**) suggesting functional complementation in mutants with *Pf*PHB proteins. EtBr is known to degrade mitochondrial DNA in yeast cells without affecting the genomic DNA (34), thereby disturbing the mitochondrial membrane potential of the cells (48). *Pf*PHBs were able to rescue the loss of mtDNA caused by EtBr in the complemented yeast cells suggesting the role of *Pf*PHB1 and *Pf*PHB2 in maintaining the integrity and homeostasis of the parasite mitochondria. Moreover, Roc-A was highlighted as targeting *Pf*PHBs by causing a possible disruption of mitochondrial membrane potential by inhibiting PHBs in the yeast cells (**Figure 9**). Thus, the observations of this study deepens the available knowledge of these *Pf*PHB proteins which can prove to beneficial in target-based drug designing studies in *P. falciparum*. Moreover, we also present Roc-A as an efficient antimalarial agent targeting PHBs in the mitochondria of the *P. falciparum*. Thereby, the inhibition of mitochondria has been suggested as a novel strategy for malaria extermination.

## ACKNOWLEDGEMENTS

We are thankful to Shiv Nadar University, RWTH Aachen University, and Jawaharlal Nehru University for providing required laboratory facilities to conduct this research. M.S. is thankful to Shiv Nadar Foundation and Deutscher Akademischer Austauschdienst (DAAD) for funding. SP is grateful for the funding support from Shiv Nadar Foundation. Funding from Intensification of Research in High Priority Areas (IRHPA) of Science and Engineering Research Board (SERB) and the National Bioscience Award from DBT for SS is acknowledged. The funders had no role in study design, analysis of results, preparation or publishing of the manuscript.

Strain BY4742 has been kindly gifted by Prof. Eric Beitz, Department of Pharmaceutical and Medicinal Chemistry, Pharmaceutical Institute, University of Kiel, Germany. PHB mutant stains SN436_Y9687::*phb1Δ* (YGR132C) and SN437_Y9687::*phb2Δ* (YGR231C) were gifted from Prof. Charles Boone, The Donnelly Centre, University of Toronto, 160 College St., Toronto ON, Canada M5S 3E1

## AUTHOR’S CONTRIBUTION

MS contributed towards experimental design, experimentation, data analysis and manuscript preparation. CJN helped in semi-quantitative PCR experiments. MM helped in yeast experiments. V conducted gametocyte inhibition experiments. KCP, SP, AR helped in experimental design and manuscript editing. GP and SS conceived the idea, experimental designing, data analysis and, approval of final draft.

